# Rclade: automated taxonomic collapsing and geological-timescale annotation of time-calibrated phylogenetic trees in R

**DOI:** 10.64898/2026.08.27.747462

**Authors:** Zichao Zeng, Yinzhao Wang

## Abstract

**Background:** Reproducible taxonomic collapsing and geological-timescale annotation of time-calibrated phylogenetic trees in R often require coordination among several packages and repeated code for label parsing, clade validation, plotting, and export. Workflow-managed analyses additionally benefit from non-interactive configuration, predictable diagnostics, and machine-readable exit status.

**Results:** We present Rclade, an R package that consolidates the multi-package coordination required for taxonomic collapsing into a streamlined, single-function interface. Rclade provides (1) custom ggproto objects (GeomPolygonStraight/GeomSegmentStraight) that bypass coord_munch() interpolation to achieve straight-edge rendering of collapsed triangles in circular layouts; (2) automatic detection and parsing of four taxonomic-label formats (GTDB, Silva, NCBI, embedded) plus user-supplied custom regex, with explicit input-validation contracts and parsing-accuracy evaluation on real and derived test sets; and (3) workflow embeddability through YAML configuration, library-mode APIs, and standard Unix exit codes. Benchmarks on synthetic and real datasets (200–10,000 synthetic tips and real reference trees up to 10,122 tips; 5 replicates at every scale under a unified fully rendered measurement protocol) show that the full-pipeline overhead is modest for interactive use (median ≈0.87 s in-session rendering and ≈8.4 s process-level wall-clock at 10,000 tips).

**Conclusions:** Rclade is a convenience layer over the ggtree/deeptime ecosystem that reduces boilerplate while adding targeted technical improvements for circular-layout rendering and format heterogeneity management.

**Availability:** Rclade is released under the MIT license. Source code, documentation, benchmark scripts, and evaluation data are archived at Zenodo (https://doi.org/10.5281/zenodo.22106523 (version 1.1.0; all versions: https://doi.org/10.5281/zenodo.22043060)), with ongoing development at https://github.com/zengzichao/Rclade.

## 1. Introduction

Phylogenetic trees are fundamental data structures in evolutionary biology. As genomic databases continue to expand—GTDB Release 232 now catalogs more than 878,000 bacterial and 22,000 archaeal genomes [1a], and its reference trees contain 189,801 bacterial (bac120) and 10,122 archaeal (ar53) tips—visualizing medium-to-large trees (hundreds to tens of thousands of tips) poses increasing challenges for visual clarity, input robustness, and workflow integration. Taxonomic collapsing, which merges branches belonging to the same taxonomic group (e.g., phylum, class) into triangular nodes, is a widely used strategy for managing tree complexity while preserving evolutionary information in both interactive [2–4] and programmatic [5] tree viewers.

However, implementing taxonomic collapsing in R requires coordinating multiple packages: ape for reading trees and computing MRCAs [6], ggtree for tree rendering [7], ggplot2 for palettes and legends [8], and deeptime for adding geological timescales [9]. A typical workflow involves dozens of lines of code, much of it boilerplate for handling different taxonomic-label formats (GTDB [10], Silva [11], NCBI [12]), managing nested branch relationships, handling multi-tree files, and validating input data quality.

Moreover, modern bioinformatics projects are often organized with workflow managers such as Snakemake or Nextflow [13,14]. These scenarios require visualization tools to be configurable through parameter files, to expose library-mode APIs for selective function calls without full pipeline execution, to support composable multi-tree output for parallel downstream processing, and to provide structured logging with standard exit codes for pipeline error capture.

We introduce Rclade, an R package that consolidates these steps into a single function call. Rclade is designed to address three practical problems that arise frequently in routine phylogenomic visualization workflows. First, ggtree’s `collapse()` produces curved triangle edges in polar-coordinate layouts due to `coord_munch()` interpolation—to our knowledge, no built-in parameter or existing extension addresses this rendering artifact as of ggtree 4.0.4; Rclade complements ggtree’s rendering capabilities through custom `GeomPolygonStraight` and `GeomSegmentStraight` ggproto objects that bypass `coord_munch()` interpolation without modifying the underlying rendering engine (§2.7 and Supporting Information, Straight-Edge Rendering section). Second, GTDB, Silva, and NCBI use mutually incompatible label formats, forcing researchers to write repetitive format-conversion code; Rclade provides a unified detection and parsing interface for four formats (plus user-supplied custom regex) with three parsing strategies for embedded labels. Third, existing visualization tools were not designed with Snakemake/Nextflow integration in mind; Rclade provides YAML configuration, standard Unix exit codes, structured logging, and HPC-safe temporary-file permissions, making tree visualization embeddable in automated workflows.

Here we present Rclade version 1.1.0, whose core contributions are: (i) straight-edge rendering of collapsed triangles in circular layouts via custom ggproto objects that bypass coord_munch() interpolation; (ii) an automated taxonomic pipeline with unified detection and parsing of four label formats plus user-supplied regex, automatic MRCA computation with nested-aware batch collapsing, and adaptive geological-timescale integration; and (iii) workflow embeddability through YAML configuration, library-mode APIs, standard Unix exit codes, and multi-tree split mode for Snakemake/Nextflow embedding.

## 2. Materials and Methods

### 2.1 Software architecture

Rclade is implemented as a standard R package (version 1.1.0) with 31 R source files organized into seven functional modules (Figure 1). The input layer (read-input.R) handles file-format detection (Newick/Nexus/BEAST XML) and basic validation. The deep validation layer (validate-deep.R, encoding.R) implements Newick syntax parsing, tree-structure checks (self-loops, multi-root, negative branch lengths, duplicate tips), sequence-file validation, and UTF-8/BOM normalization. The parsing layer (parse-taxonomy.R, taxonomy-file.R, custom-groups.R) performs taxonomic-format detection, label parsing, and external taxonomy- file merging. The MRCA and monophyly layer (compute-mrca.R, monophyly.R, special-identifiers.R) handles MRCA computation, nested-conflict detection, and special-ancestor checks. The visualization layer (plot- timetree.R, plot-timetree-pipeline.R, collapse-workflow.R, timescale.R, color-palette.R, legend-smart.R, annotate-clade.R, theme-publication.R) orchestrates rendering, batch collapsing, adaptive timescales, color-vision-deficiency-friendly palettes [15], legend layout, and branch labels. The output layer (save-timetree.R) manages multi-format export. Finally, the interface and infrastructure layer provides a CLI (cli.R), a Shiny web interface (shiny-app.R), graceful interrupt handling (interrupt.R), and structured logging (logger.R). Supporting files provide batch processing (batch.R), option and parameter plumbing (options.R, params-helpers.R), node-support label handling (support-labels.R), self-tests (selftest.R), built-in dataset documentation (data.R), and package infrastructure (logo.R, zzz.R). A workflow-integration sidebar encompasses YAML configuration, library-mode APIs, split mode, Strip Annotations, HPC-safe temporary files, CI coverage, and exit codes.

**Figure 1.**
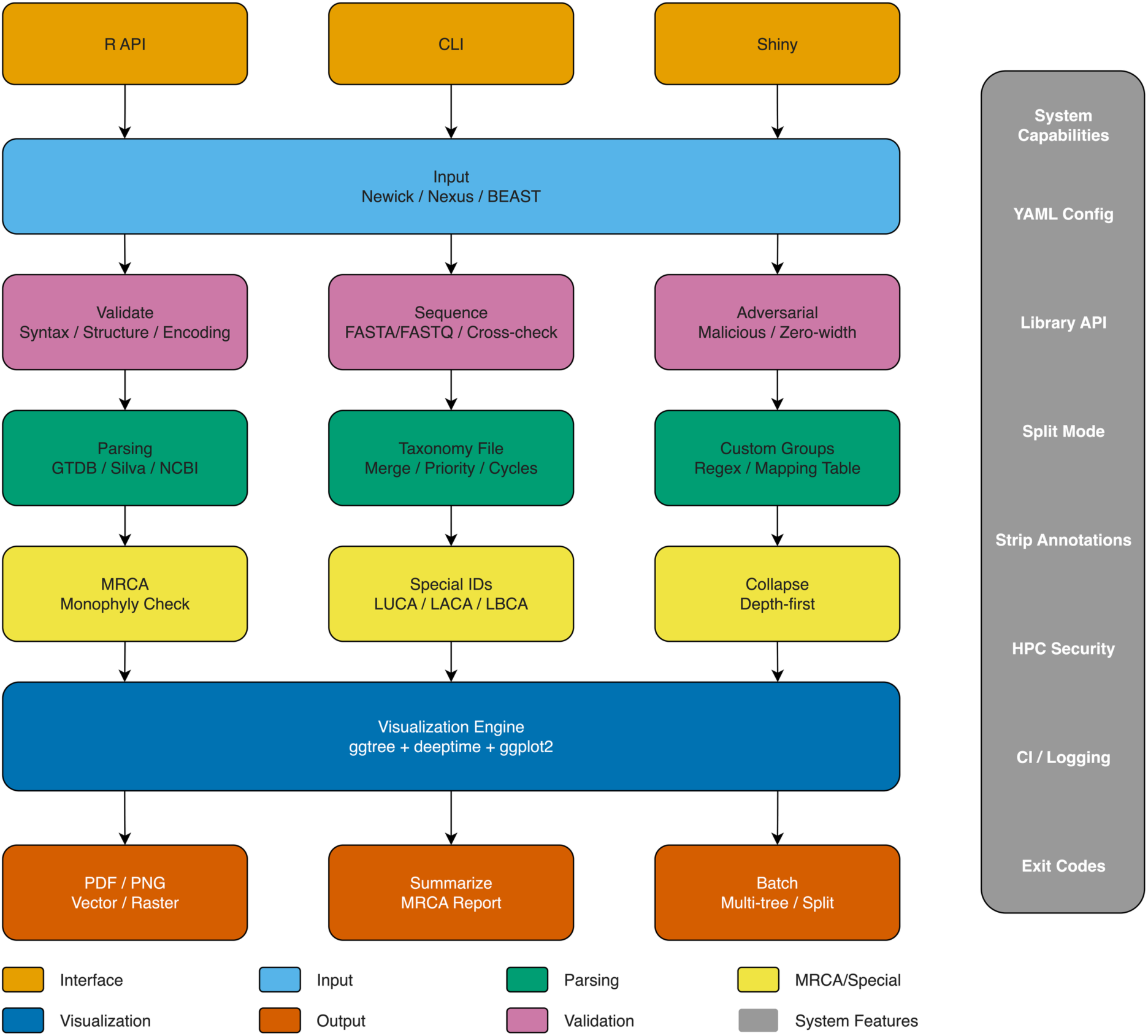
Rclade software architecture. Flowchart showing seven functional modules (seven pipeline stages) with arrows indicating data flow; a “Workflow Integration” sidebar on the right highlights additions (YAML Config, Library API, Split Mode, Strip Annotations, HPC Security, CI/Logging, Exit Codes).

**Figure 2.**
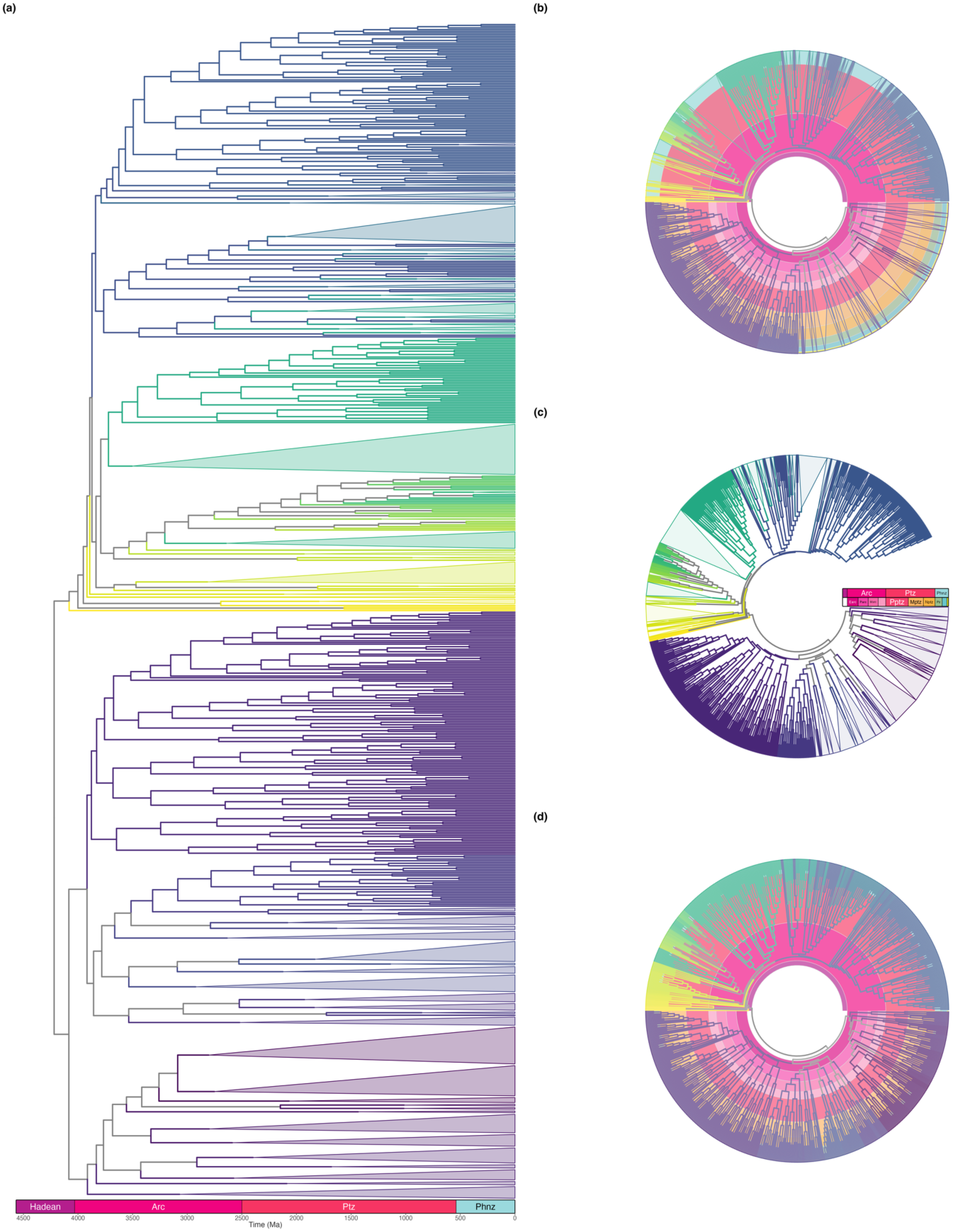
Phylum-level collapsing of a timetree. (a) Rectangular phylum-level collapsing tree with numeric time axis (Ma); the timescale shows eon-level strips only; (b) circular layout with block timescale (radial mode); (c) circular layout with background timescale (linear mode); (d) fully expanded circular tree with timescale. Branches in all panels are coloured by phylum; because this tree contains ∼125 phyla, a legend cannot fit within the main-text figure extent and is therefore omitted, so the colouring serves only to distinguish clades. Of the 125 candidate phyla parsed from this tree, 53 monophyletic groups were collapsed, 66 remained singleton tips, and 6 non-monophyletic groups were skipped (Table 4); the complete group-to-colour mapping is archived with the package (benchmark_results/gbm_phylum_color_mapping.csv). Geological-interval abbreviations on the timescale strips: Arc, Archean; Earc/Parc/Marc/Narc, Eo-/Paleo-/Meso-/Neoarchean; Pptz/Mptz/Nptz, Paleo-/Meso-/Neoproterozoic; Pz, Paleozoic; Mz, Mesozoic; Cz, Cenozoic; Ptz, Proterozoic; Phnz, Phanerozoic. (Abbreviations follow the deeptime package’s default abbreviation set rather than formal ICS conventions.) The input tree is the 700-tip archaeal timetree of Moody et al. [28] (tip labels reformatted by us to embed taxonomic lineages; topology and branch lengths unchanged). The exact reformatted input and generation script are archived with the package (inst/extdata/moody2025_gbm_laca_timetree.nwk; see Software Availability and REPRODUCIBILITY.md). Panels (b)–(d) are rendered with Rclade’s straight-edge pipeline (cf. Figure 3); all collapsed-triangle edges are straight by construction (Supporting Information, Straight-Edge Rendering section).

The package depends on eight core packages: ape (≥5.0) [6], ggtree (≥4.0.0) [7], ggplot2 (≥3.5) [8], deeptime (≥1.0) [9], rlang [16], stringr (≥1.5) [17], tidytree (≥0.4) [18], and viridisLite [15] (the lightweight implementation of the viridis package). In CI, the package is verified against the current releases of ggplot2 and ggtree (the DESCRIPTION version lower bounds are not exercised by CI) (ggtree follows the current Bioconductor release and cannot be version-pinned through BiocManager); startup diagnostics warn when major-version mismatches are detected. The development environment for this study used ggplot2 4.0.3 and ggtree 4.0.4. The custom Geom objects (`GeomPolygonStraight` / `GeomSegmentStraight`) depend on `coord$transform()` behavior and ggplot2 internal constants, which may change across ggplot2 or ggtree releases. Optional dependencies include phangorn [19] (nested detection), treeio [20] (BEAST2/IQ-TREE support), RColorBrewer [21] (additional palettes), cowplot [22]/patchwork [23] (figure composition), optparse [24] (CLI), shiny [25] (web interface), and yaml [26] (configuration files). A complete parameter reference is provided in Supplementary Table S1.

Monophyly check default behavior: In the plot_timetree() pipeline, compute_mrca_map() enables monophyly checking by default (check_monophyly = TRUE), with non-monophyletic groups producing a warning and being omitted from the collapse plan by default (warn-and-skip; strict = FALSE); singleton groups are likewise retained as individual tips rather than collapsed. Users can escalate non-monophyletic groups from warning to error via the --strict CLI option or by setting strict = TRUE directly. This default ensures predictable behavior in automated pipelines—runs will not be aborted due to monophyly issues by default, but users are notified via warnings and the completion summary reports per-group status (total/collapsed/singleton/skipped), since a successful non-strict run may represent a partial collapse.

### 2.2 Configuration-driven workflows

Rclade 1.0.1 introduces YAML configuration-file support (`--config`). Parameter priority is **CLI explicit arguments > config file > built-in defaults**. Config files use long parameter names as keys (e.g., `rank`, `color_palette`, `log_level`); booleans use native YAML booleans, and lists or maps use comma-separated strings for CLI compatibility.

Configuration-driven workflows are especially useful for providing unified default styles across all samples in a Snakemake rule, reusing visualization parameters across papers and projects, and quickly generating results for different ranks by overriding a single CLI parameter (Figure S1).

### 2.3 Library-mode API

In addition to the high-level end-user interface, Rclade exports 26 functions, among which two are designated as the stable low-level API for direct use by external workflows: `Rclade::read_tree_auto(filepath, tree_index, multi_tree_mode)` automatically detects file format, handles multi-tree files, and returns a `phylo` or `multiPhylo` object; `Rclade::parse_taxonomy(labels, rank, format, …)` parses taxonomic labels into a standard data.frame, supporting GTDB, Silva, NCBI, embedded, and custom_regex formats.

These functions allow Snakemake/Nextflow pipelines to complete format detection, label parsing, and tree reading without triggering the full plot_timetree() rendering, and then pass results to downstream statistical analysis or custom visualization (Figure S2).

### 2.4 Multi-tree split mode

Multi-tree files (e.g., BEAST posterior `.trees`) traditionally require users to select a specific tree or use the `all` mode for batch output. 1.0.1 adds `--multi_tree_mode split`, which has output behavior similar to `all` (all trees are processed) but explicitly conveys the “split output per tree” semantics, so that external workflows can dispatch one downstream task per tree; split mode itself processes trees sequentially and does not schedule parallel jobs (Figure S3).

### 2.5 Engineering practices

Rclade implements engineering conventions to support deployment in automated pipelines: standard Unix exit codes (0/1/2/3/130) with a stable SIGINT wrapper (`inst/bin/rclade`)—on HPC schedulers, job preemption or user cancellation must surface as exit code 130 rather than a generic failure so that downstream workflow rules can distinguish interrupts from genuine errors; HPC-safe temporary-file permissions (0600/0700); structured logging with module tags; CI coverage thresholds (85%); and a `--strip_annotations` option for reducing output size (Figure S6). These are standard practices for pipeline-embeddable tools and are detailed in the Supplementary Methods.

### 2.6 Retained core methods (summary)

Rclade’s taxonomic-format detection, MRCA/monophyly computation, adaptive timescale, in-depth input validation, and graceful interrupt handling have been iteratively developed and matured. 1.0.1 preserves these methods while enhancing integrability through the engineering improvements above. This section briefly reviews the key mechanisms.

#### 2.6.1 Taxonomic-format detection

`detect_taxonomy_format()` automatically detects GTDB, embedded, Silva, and NCBI formats based on prefix-matching scores. For embedded formats, `reverse` (default), `greedy`, and `segment` delimiter-matching strategies are provided. Users may also supply custom regex patterns via `taxonomy_format = "custom_regex"` for non-standard label schemes, in which case format detection is bypassed. The detection algorithm samples the first 100 tip labels and applies the following rules sequentially:

GTDB/embedded use the “clear majority” rule (score > 0.5 and at least 0.1 above the 0.5 tie line, i.e. score ≥ 0.6); NCBI/Silva use a lower “dominance” threshold (score > 0.3) because semicolon-delimited formats have higher prefix diversity, so individual prefix match rates are naturally lower. The GTDB rule additionally requires a semicolon majority: accession-prefixed embedded labels that use double-underscore rank separators (e.g., `GCA_xxx_d` `Archaea_p` `Nanoarchaeota`, as found in real timetree exports) also satisfy the `[dpcofgsk]` pattern but contain no semicolons, and would otherwise be misclassified as GTDB (see §3.5 for this real-world case).

The 0.6 clear-majority threshold requires that a format’s prefix-match score is strictly above 0.5 and at least 0.1 away from the 0.5 tie line (i.e. score ≥ 0.6): when no format reaches a clear majority, the evidence is too ambiguous, and conservative fallback to “unknown” is safer than forcing a potentially incorrect classification. This threshold’s behavior was characterized under increasing random-label noise on GTDB-style labels (Figure S4): under that idealized construction the prefix-match score equals 1 − f, so detection deterministically succeeds while noise ≤ 40% and falls back to “unknown” beyond it; Figure S4 illustrates this decision rule rather than an empirical accuracy estimate.

#### 2.6.2 Performance benchmarks

Rclade’s runtime and memory on synthetic datasets were measured with in-session bench::mark medians [27] (5 replicates per configuration) for rendering times, and /usr/bin/time -l (5 independent process runs per configuration) for process-level wall-clock and peak memory. Within the tested range the median runtime grew approximately linearly with the number of tips, reaching ≈0.87 s (in-session rendering) / ≈8.4 s (process-level wall-clock, including R startup and package loading) at 10,000 tips; peak resident memory at 10,000 tips was ≈3.6 GB in the default configuration (≈679 MB with the optional low_memory garbage-collection mode) (Figure S5, Table 3).

**Table 1.** Taxonomic-format detection: decision rules and thresholds.

| Format | Decision rule | Threshold |
| --- | --- | --- |
| GTDB | Fraction of labels containing [dpcofgsk]__ prefixes (clear majority) **and** semicolon-delimiter fraction > 0.5 | score ≥ 0.6 |
| embedded | Fraction of labels matching _[dpcofgsk]_ pattern (clear majority) | score ≥ 0.6 |
| NCBI / Silva | First require semicolon-delimiter fraction > 0.5, then compare NCBI prefix score vs Silva prefix score; the dominant one must exceed threshold | dominant score > 0.3 |
| unknown | Fallback when tied or both below threshold | — |

**Table 2.** Time complexity of each pipeline stage.

| Stage | Complexity | Notes |
| --- | --- | --- |
| Label parsing (parse_taxonomy) | O(n) | One regex match per label |
| Format detection (detect_taxonomy_format) | O(min(n, 100)) | Samples first 100 labels |
| MRCA computation (compute_mrca_map) | O(k · n) | One ape::extract.clade() call per group |
| Nested-conflict detection (validate_collapse_plan) | O(k<sup>2</sup>) | Pairwise descendant-intersection checks |
| Collapse rendering (collapse_by_groups) | O(k) | One collapse() + layer overlay per group |
| Timescale integration | O(1) | Independent of tree size |

**Table 3.** Rclade full-pipeline overhead characterization on synthetic datasets (unified measurement protocol).

| Tips | Groups (k) | Rclade in-session (s)† | Core collapsing only† (s) | Overhead factor‡ | Process wall-clock (s)§ | Rclade peak memory (MB)¶ |
| --- | --- | --- | --- | --- | --- | --- |
| 200 | 14 | 0.36 | 0.020 | 17.8× | 2.01 | 522 |
| 1,000 | 31 | 0.44 | 0.021 | 21.0× | 2.31 | 570 |
| 5,000 | 70 | 0.55 | 0.027 | 20.1× | 4.08 | 1,421 |
| 10,000 | 100 | 0.87 | 0.034 | 25.3× | 8.37 | 3,653 |

† In-session columns are bench::mark medians in a warm R session (5 replicates per scale; scripts/benchmark\_synthetic.R), excluding process startup and package loading. The core-collapsing-only baseline performs MRCA computation, scaleClade(vertical\_only = TRUE), and ggtree::collapse() (mode = “mixed”), excluding Rclade’s automation features (format detection, label parsing, monophyly checking, timescale integration, legend layout, input validation, logging, and palette generation); Rclade forces mode = “max” in circular layouts, so this comparison quantifies the cost of automation features rather than a like-for-like speed comparison. Replicates are right-skewed at large n (at 10,000 tips: mean 2.33 s, sd 2.50 s, median 0.93 s; individual measurements in benchmark\_results/benchmark\_synthetic\_rendered.csv), so medians are reported throughout; the skew reflects OS-level background load on the laptop test platform.
‡ Overhead factor = Rclade in-session / core collapsing only (in-session). Values > 1 indicate the additional cost of automation features.
§ Process wall-clock from the median of 5 independent /usr/bin/time runs per configuration (scripts/run\_process\_level.sh), including R startup, package loading, I/O, and computation.
¶ Memory values are peak RSS from /usr/bin/time -l (median of 5 independent runs per configuration), subject to macOS memory compression and system caching. Measurement-level note: Table 3 reports two complementary series under one unified protocol—in-session rendering medians (bench::mark, warm session; also plotted in Figure S5) and process-level wall-clock (/usr/bin/time, including R startup and package loading). The process-level series adds an approximately constant startup overhead to both pipelines (≈1.8 s for Rclade vs ≈1.3 s for the baseline at n = 200), compressing the process-level overhead ratio to ≈1.5–2.3× versus the 18–25× in-session ratio; the two series answer different questions (deployment cost vs incremental render cost) and are not expected to coincide. Raw measurements are provided in benchmark\_results/ (benchmark\_synthetic\_rendered.csv, process\_level\_metrics.csv, benchmark\_split\_cost\_1000.csv).

#### 2.6.3 Computational complexity

The time complexity of each pipeline stage is as follows (n pa = number of tips, k = number of collapsing groups):

The conservative overall bound is O(n·k + k²) plus renderer-dependent layer costs; where k ≪ n and k is bounded this reduces to approximately linear, consistent with the observed medians over the tested 200–10,000-tip range. Note that the O(k) collapse-rendering stage creates two ggplot2 layers per group color (fill + border; colors are passed as non-aesthetic parameters to avoid scale conflicts), so at order rank the layer count reaches the hundreds and dominates the rendering-side constant factor (see §4.3, Limitation 4).

### 2.7 Straight-edge rendering of collapsed triangles in circular layout Problem

In circular (fan) layout, taxonomic collapsing compresses monophyletic clades into triangular nodes to reduce visual clutter; however, ggtree’s collapse() produces collapsed triangles with curved edges rather than the expected straight lines, which can reduce the aesthetic quality and readability of publication-ready figures, especially when collapsed triangles are large.

#### Limitations of existing tools

The artifact originates from ggplot2’s internal call to `coord_munch()` under `coord_polar()`, which interpolates polygon edges into segments to simulate polar curves, bending originally straight edges. To our knowledge, as of ggtree 4.0.4, `collapse()` provides no parameter to disable the interpolation, and no existing ggplot2/ggtree extension addresses this specific rendering case (searched alternatives are listed in the Supporting Information, “Searched alternatives” section). More critically, even with straight-edge triangles, the MRCA vertex lies outside the bounding box of the tip nodes and is truncated under standard panel clipping, so the apex of the top triangle is missing—an integrity issue that existing rendering pipelines generally overlook.

#### Method

Rclade complements ggtree’s rendering capabilities at the correct abstraction layer (custom ggproto geometric objects) rather than modifying the core logic of ggtree/ggplot2: (1) GeomPolygonStraight and GeomSegmentStraight transform only the vertex coordinates via coord$transform() into polar coordinates and draw straight-edge primitives in npc space using grid’s polygonGrob/segmentsGrob, thereby completely bypassing coord_munch() interpolation; (2) collapsed-triangle vertices are pre-computed before calling collapse(), and the fixed 0.5 offset of get_clade_position() is corrected using the actual tip spacing so that triangle size is proportional to the number of tips in the collapsed clade; (3) circular layout forces “max” mode so that triangle vertices align precisely with the outer edge of the tree.

#### Implementation and tested behavior

Fill and border colors are passed through non-aesthetic parameters to avoid conflicts with the `scale_fill_manual` legend scale (eliminating the “No shared levels found” warning). To address the vertex over-run issue, the rendering layer disables panel clipping (`coord_cartesian(clip = "off")`), so that in all tested configurations the MRCA vertex is displayed completely outside the panel boundary; together with the tip-spacing adaptive vertex correction above, in tested configurations this prevents vertices from being censored to NA by the out-of-bounds handler or clipped. This technique does not modify the core rendering logic of ggtree/ggplot2 but extends their functionality only through custom ggproto layers. A known limitation of this approach is that the custom Geom objects depend on `coord$transform()` behaviour and ggplot2 internal constants (e.g., `.pt`), which are not part of the public API and may change across ggplot2/ggtree releases; version- compatibility guards are implemented at package load time (see §4.3). The rendering-pipeline principles and effect comparison are presented in the Results section (§3.4, Figure 3); the full technical principles, geometric derivations, code implementation, and limitations are provided in the Supporting Information (Straight-Edge Rendering section).

**Figure 3.**
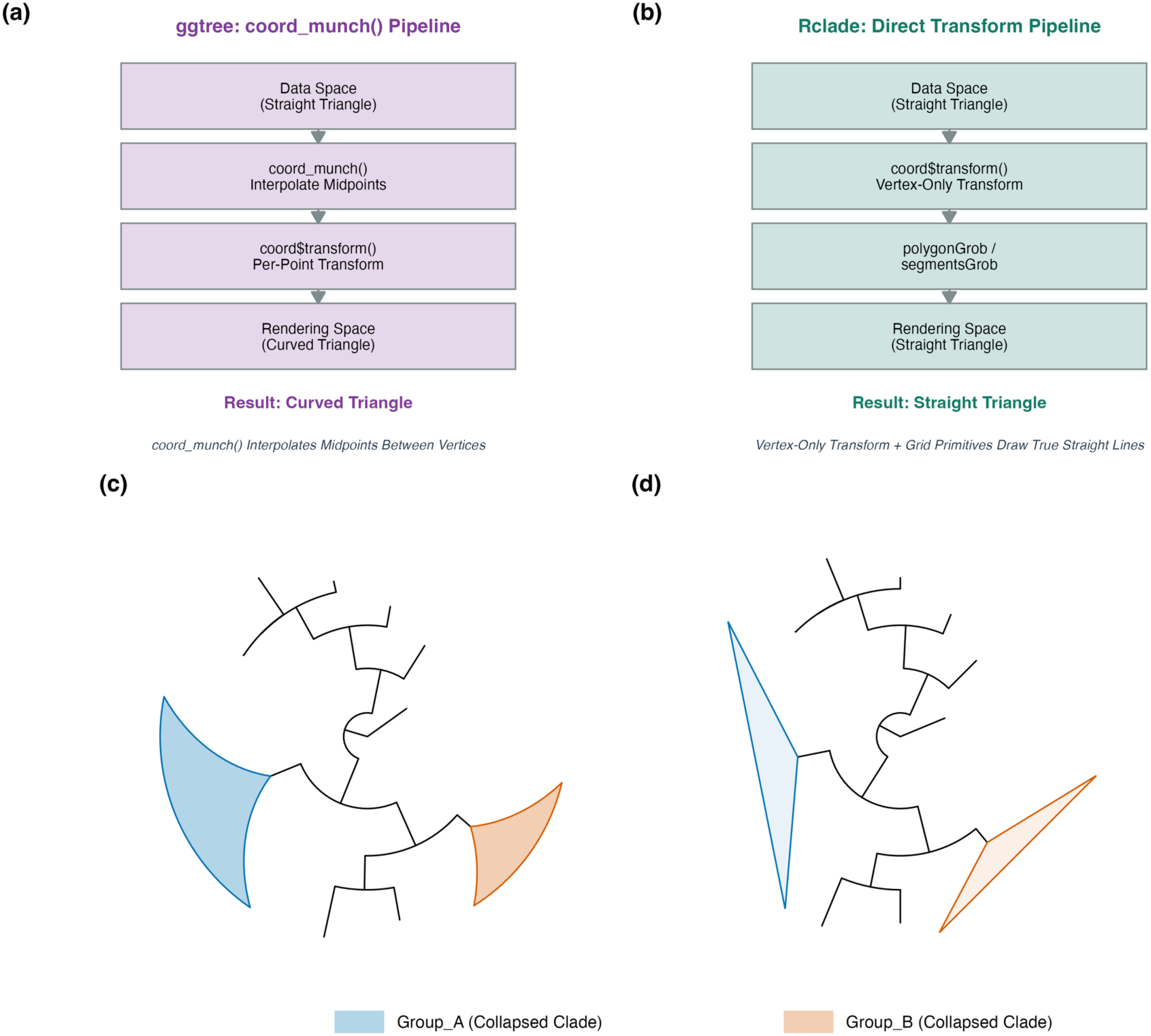
Straight-edge rendering technology for collapsed triangles in circular layout. (a) ggtree: coord_munch() rendering-pipeline schematic, resulting in curved triangles. (b) Rclade: direct-transform rendering-pipeline schematic, resulting in straight triangles. (c) Actual rendering of ggtree’s native collapse() on a 15-tip synthetic small tree, with curved collapsed-triangle edges. (d) Actual rendering of Rclade’s straight-edge rendering on the same small tree, with straight collapsed-triangle edges. The two collapsed clades use consistent colors in (c) and (d).

## 3. Results

### 3.1 Workflow-integration validation

#### 3.1.1 YAML config + CLI override

Using `v4_config.yaml` as the default (`rank = none`, `layout = rectangular`, `color_palette = viridis`) and overriding only the `-r` parameter via the CLI, we generated phylum-, class-, and order-level visualizations in three commands (Figure S1). This pattern is particularly useful in Snakemake rules: the rule body

only needs to specify input/output and the parameters to override, while styling is controlled uniformly by the config file.

#### 3.1.2 Library-mode API

In an independent R session, we verified that `read_tree_auto()` correctly reads single- and multi-tree Newick files, `parse_taxonomy()` parses GTDB/Silva/NCBI labels into a uniform data.frame, and `detect_taxonomy_format()` returns “GTDB” for the built-in `example_tree`. These functions do not depend on ggplot2/ggtree rendering and can run on headless servers, making them suitable as parsing steps in pipelines.

#### 3.1.3 Multi-tree split mode

Running `--multi_tree_mode split` on a `.trees` file containing 3 synthetic trees produced 3 independent output files with automatically appended index suffixes. Internally, `split` mode shares the same code path as `all` mode (both return a `multiPhylo` object and iterate over each tree), but `split` explicitly conveys the “split output per tree” semantics, providing a clearer interface contract for downstream pipelines. Structured per-tree metadata output (JSON/YAML) is planned for future versions (§4.4).

#### 3.1.4 Web interface

`run_rclade_shiny()` launches a Shiny-based web interface that wraps `plot_timetree()` with interactive parameter controls (tree upload, rank/layout/palette selection, live preview, and export). Interface startup and control wiring are covered by the package test suite (`test-shiny.R`), ensuring the GUI remains functional across dependency updates.

### 3.2 Performance and resources

We profiled Rclade’s full-pipeline overhead on synthetic datasets to characterize how much time the automation features (format detection, monophyly checking, timescale integration, input validation, legend layout) add on top of the core collapsing operation.

#### Test platform

All benchmarks in this study were conducted on a single personal laptop (MacBook Pro, Mac17,2; Apple M5; 32 GB unified memory; macOS 26.6; micromamba environment R 4.5.3). All in-session scales were measured with 5 replicates; process-level wall-clock and peak RSS were measured with 5 independent process runs per configuration. The absolute time and memory values reported here should be considered reference data for this specific hardware configuration rather than universal performance guarantees; because the measured workflow is predominantly single-threaded, additional CPU cores are not expected to speed up a single run; absolute performance may differ with per-core speed, memory capacity, operating system, and dependency versions.

Within the tested 200–10,000-tip range the median fully rendered runtime was approximately linear (≈0.87 s at 10,000 tips); per-run times are right-skewed (max ≈5.7 s at 10,000 tips), so individual runs can be slower.

The 18–25× in-session overhead factor reflects Rclade’s automation features—format detection, monophyly checking, timescale integration, input validation, legend layout, and palette generation—layered on a fast core collapsing operation. To localize this cost, split-stage timing at n = 1,000 (3 replicates, proc.time protocol including timescale and legend; scripts/benchmark_split_cost.R) attributed 10.4% (0.12 ± 0.00 s) to format detection and parsing, 2.0% (0.02 ± 0.00 s) to MRCA + monophyly, and 28.1% (0.33 ± 0.08 s) to collapsing and rendering excluding timescale and legend; the remaining ∼59.5% is not attributed to individual stages because the split-stage protocol differs from the end-to-end protocol of Table 3. Peak resident memory (low_memory mode) grew gently with tree size to 679 MB at 10,000 tips and was dominated by R-session and dependency overhead, while CPU usage approached 100% on a single core, reflecting R’s single-threaded nature.

On the 10,122-tip archaeal reference tree, Rclade completed phylum-, class-, and order-level collapsing in 4.3–6.4 s (in-session medians; 5.6–9.3 s process wall-clock) with a peak memory of ∼613–666 MB. The real-data in-session times exceed the synthetic 10,000-tip case (0.93 s) despite comparable tip counts; the difference is attributable to the real-data workload carrying accession-only tip labels resolved through the external taxonomy- file interface, denser internal node annotations, and higher collapsed-group counts (up to 179 orders versus 100 synthetic groups), which increase MRCA computation, legend layout, and per-color layer overhead.

### 3.3 Application examples

#### Example 1: Phylum-level collapsing of a timetree

Using the 700-tip archaeal timetree published by Moody et al. [28] (the exact reformatted input used here is archived with the package at inst/extdata/moody2025_gbm_laca_timetree.nwk; see Software Availability and REPRODUCIBILITY.md) for phylum-level collapsing, demonstrating Rclade’s rendering capabilities across different layouts and timescale modes. The combined figure (Figure 2) uses a 2-column layout: the left column shows rectangular phylum-level collapsing; the right column contains three panels from top to bottom—circular phylum-level collapsing with block timescale (radial mode), circular phylum-level collapsing with background timescale (linear mode), and a fully expanded circular tree with timescale. This example demonstrates Rclade’s ability to render the same tree consistently with different layouts and timescale modes, all generated through single function calls.

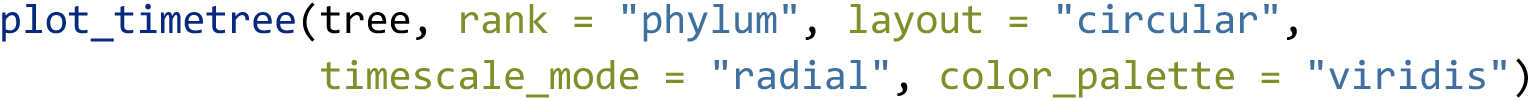

**Table 4.** Real-dataset benchmarks.†.

| Dataset | Tips | Rank | Groups<br>(parsed/collapsed) | Displayed leaves<br>after collapse | In-session<br>median (s)‡ | Process wall-clock (s)§ | Peak memory<br>(MB)¶ |
| --- | --- | --- | --- | --- | --- | --- | --- |
| ar53 (GTDB R232) | 10,122 | phylum | 25/23 | 25 | 4.24 | 5.64 | 666 |
| ar53 (GTDB R232) | 10,122 | class | 71/62 | 71 | 5.02 | 6.74 | 666 |
| ar53 (GTDB R232) | 10,122 | order | 179/142 | 179 | 7.31 | 9.29 | 668 |
| GBM LACA timetree | 700 | phylum | 125/53 | 437 | 0.57 | — | — |

† The ar53 rows show the same GTDB archaeal reference tree tested at different ranks; in-session values are bench::mark medians of 5 replicates (scripts in the Zenodo archive); process-level values are medians of 5 independent /usr/bin/time runs (re-measured 2026-08-26 under the Rclade 1.1.0 unified protocol). “Groups (parsed/collapsed)” reports candidate groups parsed and clades actually collapsed (valid, monophyletic MRCAs); singleton and skipped non-monophyletic groups are excluded from the collapsed count. “Displayed leaves after collapse” counts visible true tips plus collapsed pseudo-leaf clades (§4.3, Limitation 6). Collapsed-group counts differ from earlier internal runs because the taxonomy TSV is a newer R232 extraction and the format-detection rule was corrected (§3.5). The GBM LACA row uses the 700-tip archaeal timetree of Moody et al. [28] (embedded double-underscore labels parsed via custom\_regex, §3.5; only in-session timing recorded). Memory values are medians of peak RSS across the 5 process runs.
‡ In-session bench::mark median, as in Table 3 †.
§ Process wall-clock, as in Table 3 §.

#### Example 2: Config-file batch processing

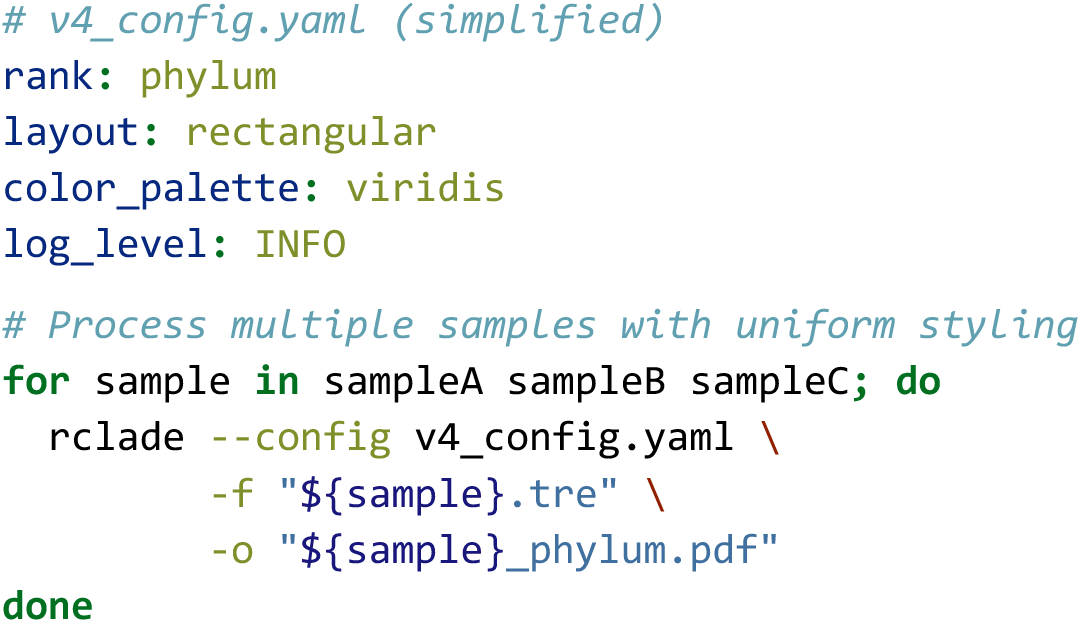

#### Example 3: Snakemake-style integration

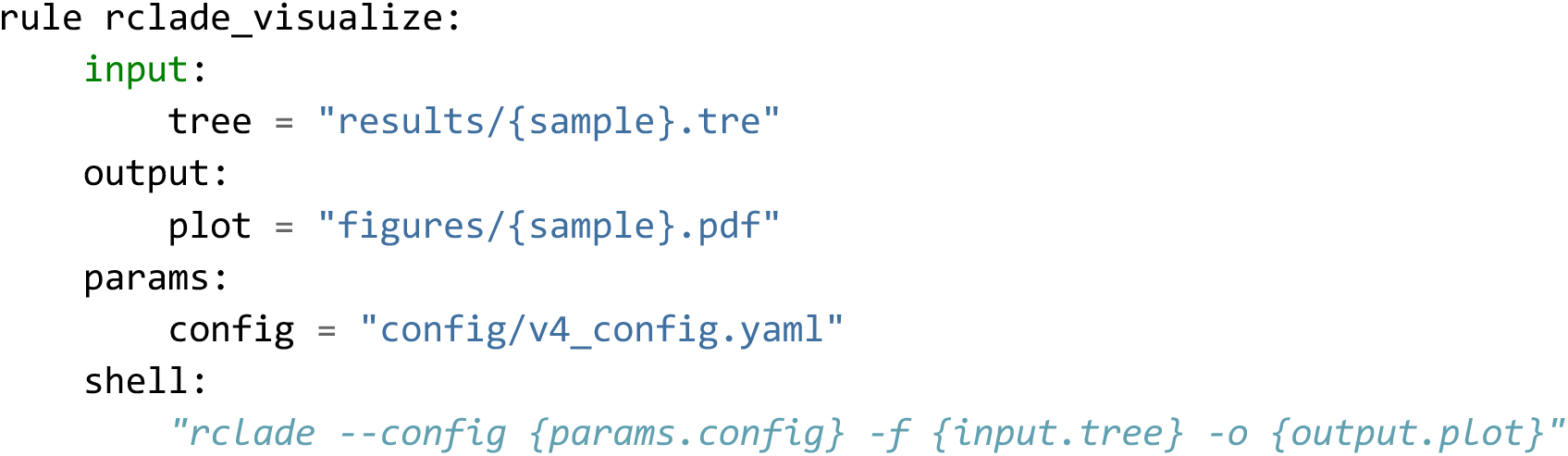

### 3.4 Straight-edge rendering effect in circular layout

To validate the effectiveness of the custom `GeomPolygonStraight` and `GeomSegmentStraight` ggproto objects, we conducted a comparative experiment using a synthetic small tree (set.seed(42) + rtree(15)) with phylum-level collapsing in circular layout. We note that this small tree (15 tips) was chosen for visual clarity in demonstrating the rendering-pipeline differences; the effect is equally present and arguably more impactful on larger trees where collapsed triangles span wider angular ranges (see GTDB ar53 examples in Table 4). Figure 3 uses a 2×2 layout to present four aspects: (a) the ggtree coord_munch() rendering-pipeline schematic, illustrating the mechanism by which midpoint interpolation causes triangle edges to curve; (b) the Rclade direct-transform rendering-pipeline schematic, illustrating the strategy of transforming only vertex coordinates to keep edges straight; (c) the actual rendering of ggtree’s native collapse() on the 15-tip small tree, where collapsed triangle edges are curved; (d) the actual rendering of Rclade’s straight-edge rendering on the same small tree, where collapsed triangle edges remain straight. The two collapsed clades (Group_A, Group_B) use consistent colors in (c) and (d) to facilitate direct comparison of curved versus straight effects.

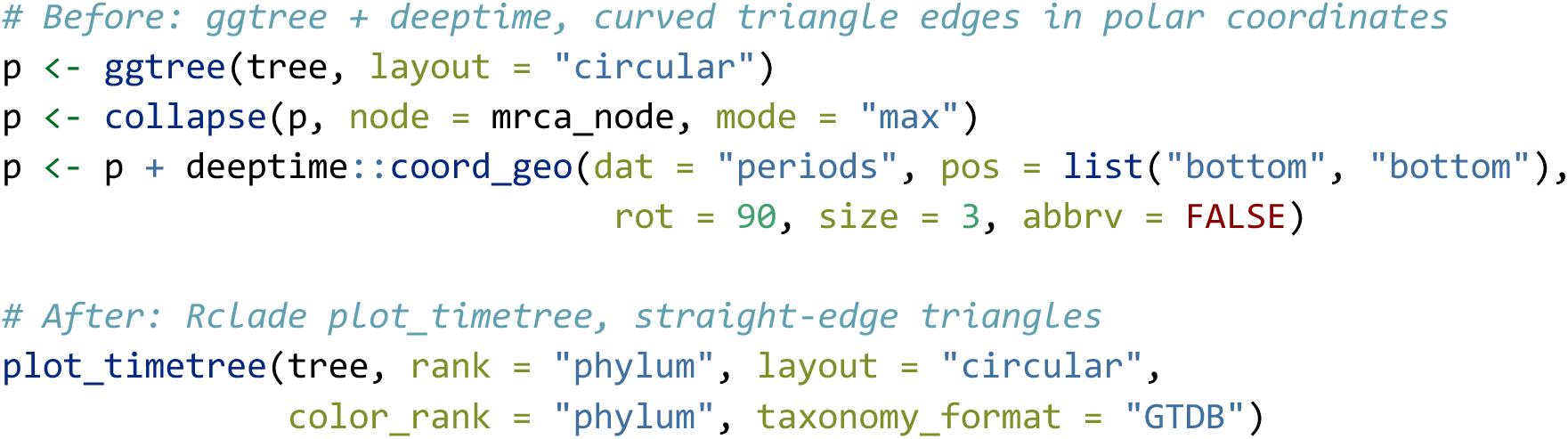

To move beyond this qualitative comparison, we measured the maximum perpendicular deviation of each collapsed-triangle edge from its ideal straight chord in transformed (npc) space, on the identical Figure 3 tree (`scripts/measure_straightness.R`). Under the ggtree pipeline, `coord_munch()` interpolates each triangle into 57–74 transformed points and deviates from the straight chord by up to 0.050–0.086 npc (mean 0.027–0.051 npc), corresponding to ≈5.5–9.5 mm on a 110 mm rendered panel—clearly visible curvature. Under Rclade’s vertex-only pipeline, exactly 3 transformed vertices are emitted per triangle and the measured deviation is 0 by construction (benchmark_results/straightness_deviation.csv). This quantitative gap confirms that the rendering difference is structural rather than cosmetic, and it grows with triangle angular span on larger trees.

### 3.5 Taxonomic label parsing accuracy

Parsing accuracy was first verified on the built-in `example_tree` (50 tips, GTDB-style labels) with 100% correctness at both phylum and class ranks (Figure S7; validation data available from the GitHub repository). This result is expected for well-formed GTDB labels where prefix-matching is deterministic—it confirms that the regex-based parsing logic is correctly implemented for the canonical case. We then extended the evaluation to real datasets with ground-truth comparison at every rank (the GTDB evaluation script and results are included in the Zenodo archive).

#### GTDB (10,122 real database labels)

On the complete GTDB R232 ar53 taxonomy table (including placeholders and candidatus-style names), parse_taxonomy() achieved a 100% non-NA parse rate **and** 100% exact-match agreement against the source fields at every rank from domain to species, confirming that the GTDB parser handles real database strings rather than only synthetic ones.

#### Embedded format on a real 700-tip timetree

We evaluated the published archaeal timetree of Moody et al. [28] (700 genome tips; BEAST posterior export with 95% CI node annotations). The tree topology, branch lengths, and node annotations are unchanged from the original publication; the tip labels were reformatted by us to embed accession-prefixed, double-underscore GTDB-style lineages; the exact reformatted file is archived with the package (inst/extdata/moody2025_gbm_laca_timetree.nwk), and all numbers below reproduce from it (benchmark_results/embedded_evaluation_moody700.csv) with an extra “superphylum” field and underscore- bearing genus names (e.g., `GCA_000387965_d` `Archaea_superphylum` `DPANN_p` `Nanoarchaeota_…_g` `Candidatus_Nan obsidianus`). This label scheme exposed two real-world issues that were subsequently fixed: (i) format detection originally misclassified these labels as GTDB (the `[dpcofgsk]` pattern matched the double underscores) and parsed 0% of phyla; detection now requires a semicolon majority for the GTDB rule, and correctly returns “embedded” for this tree; (ii) the embedded parsers now tolerate double-underscore rank separators. After the fix, the three `delimiter_mode` strategies behaved as designed: `reverse` and `greedy` conservatively returned NA for labels they could not resolve unambiguously (covering 90.0–99.6% of labels at phylum–genus ranks, with 100% exact-match accuracy among covered labels—no misassignments; Candidatus-style underscore genus names appear as ungrouped; the domain column is unresolved (0% coverage) by all built-in strategies because this scheme inserts an unmodeled superphylum field directly after the domain token), while `segment` covered 100% of labels with 90.0–100% per-rank accuracy at phylum–genus (genus 100%, including underscore-bearing names); for the domain column, `segment` captures the adjacent unmodeled superphylum field instead of the domain name (0% exact match for that single column). User-supplied `custom_regex` patterns achieved 100% non-NA rate and 100% exact-match accuracy at all six ranks, motivating the documented guidance: non-standard embedded schemes with extra ranks should use `custom_regex`, and labels with embedded underscores should prefer `delimiter_mode = "segment"` over the conservative default.

#### Silva and NCBI (position-based parsers)

To evaluate the two position-based parsers on real taxonomic content, the 10,122 real GTDB R232 archaeal lineages were reformatted to each database’s conventions (unprefixed semicolon-delimited for Silva; “cellular organisms;”-prefixed for NCBI) and re-parsed. Both parsers achieved 100% non-NA rate and 100% exact-match agreement at all seven ranks, confirming correct positional mapping for standard 7-rank lineages (the reformatting step is disclosed; the taxonomic content is real). Because reformatting cannot capture every idiosyncrasy of native Silva/NCBI strings (e.g., “uncultured” placeholders, parenthesised synonyms), users are advised to verify parsed output for such labels; targeted evaluation on native-format samples is planned as future work.

#### NCBI positional rank shift (quantified)

The position-based NCBI mapping is known to shift ranks for non-standard lineage depths (§4.3, Limitation 1). We quantified this on an 8-token virus-style lineage: after removing the leading skip-prefix “Viruses”, the remaining 7 tokens (realm→species) map positionally: the seven positional columns receive tokens shifted by 1–2 true ranks (the phylum and class columns shift by one rank; the order, family, genus, and species columns shift by two), e.g., the species column receives the family-level token (`Filoviridae`) while the true genus and species tokens are dropped entirely (benchmark_results/ncbi_rank_shift_example.csv). Users analyzing viruses or deep environmental lineages should therefore use GTDB/Silva formats or custom regex, and Rclade emits a warning whenever NCBI positional mapping is used.

Format-detection robustness under progressively degraded inputs (mixed random labels) is evaluated separately in Figure S4 (GTDB-style labels), which tests the system’s degradation under noise rather than parsing correctness per se.

## 4. Discussion

### 4.1 Positioning and contribution

We want to be explicit about Rclade’s positioning:

### What Rclade is

a convenience wrapper that integrates existing ggtree and deeptime functionality; a format- handling layer that manages heterogeneity among taxonomic databases; a robustness layer providing structured input validation and informative error reporting; a workflow standardizer that supports reproducible tree visualization; a visualization component embeddable in production pipelines such as Snakemake/Nextflow; and a documented package with an automated test suite and multiple interfaces.

Rclade does not implement new phylogenetic algorithms, statistical models, or inference methods, and is not intended as a replacement for iTOL, FigTree, or ggtree—it complements the ggtree/deeptime ecosystem. 1.0.1 extends the contribution from “reducing code volume” to “reducing pipeline-integration cost”. The code-volume reduction is concrete: the equivalent manual workflow (Supporting Information, Baseline Script section; `scripts/baseline_manual_workflow.R`) requires ∼100 lines coordinating five packages, whereas Rclade completes the same task with a single `plot_timetree()` call.

### 4.2 Comparison with existing tools

Rclade occupies a specific position in the phylogenetic-visualization landscape:

Runtime comparisons against iTOL and FigTree are not feasible in a scriptable, like-for-like manner because both are interactive tools without equivalent batch collapsing pipelines; we therefore benchmark against a minimal ggtree-based script implementing the same collapsing semantics (Table 3), and compare functionality qualitatively (Table 5). Rclade does not compete with iTOL or FigTree in interactive exploration. It targets users who need reproducible, programmatic tree visualization within the R ecosystem. Related packages in the broader ecosystem include ggtreeExtra [31] for additional annotation layers, TreeViewer [32] for interactive exploration with publication-quality export, phytools [33] for comparative-method visualizations, and treeio [20] for multi-format tree I/O; Rclade complements rather than replaces these tools by providing automation at the collapsing and timescale-integration layer.

**Table 5.** Feature comparison of Rclade with existing phylogenetic-visualization tools.

| Feature | iTOL [2,3] | FigTree [4] | ggtree [7,29,30] | Rclade |
| --- | --- | --- | --- | --- |
| Interface | Web | GUI | R API | R API + CLI + Shiny |
| Taxonomic collapsing | Manual | Manual | Semi-automatic† | Automatic |
| Geological timescale | No | No | Via deeptime | Integrated |
| Format auto-detection | No | No | No | Yes (4 formats + custom regex) |
| Circular straight-edge | Not addressed | Not addressed | No | Yes |
| Batch processing | API‡ | Limited§ | Manual | Built-in |
| Config-driven workflow | No | No | No | YAML |
| Target use case | Interactive exploration | Interactive exploration | Programmatic R workflows | Automated pipelines |
| Ecosystem compatibility | Standalone | Standalone | Full R/Bioc | Full R/Bioc |

† ggtree provides collapse(node=N) for individual-node collapsing and groupClade()/groupOTU() for clade-based annotation [7]; the ggtree object framework further supports mapping and visualizing associated data on phylogenies [29,30]. Rclade adds automatic taxonomy parsing and batch MRCA computation on top of these primitives.
‡ iTOL v5/v6 supports batch dataset uploads through its web service (third-party scripting wrappers exist), but taxonomic annotation requires manually prepared dataset files; it does not support automatic label parsing or batch collapsing.
§ FigTree supports batch image export via command-line arguments but does not support parameterized programmatic control.

### 4.3 Limitations

#### Several limitations should be noted

Limitation 1. NCBI parsing uses position-based rank mapping, which may produce systematic rank shifts for non-standard lineages such as viruses: on an 8-token virus-style lineage, parsed columns shift by 1–2 true ranks and the true genus/species tokens are dropped (benchmark_results/ncbi_rank_shift_example.csv). Users analyzing viruses or deep environmental lineages should use GTDB or Silva formats, or supply custom regex patterns; Rclade warns whenever NCBI positional mapping is used. Relatedly, native-format SILVA/NCBI validation remains limited to derived format-conversion test sets based on real GTDB taxonomic content, and the 700-tip embedded evaluation uses author-reformatted labels of a published topology (§3.5).

Limitation 2. Taxonomic-format detection is heuristic rather than statistically optimal and may fail for heavily modified label formats; the taxonomy_format parameter allows manual override, and the 60% clear-majority threshold conservatively returns “unknown” near the boundary (Figure S4). Embedded labels carrying unmodeled intermediate ranks (e.g., a “superphylum” field between domain and phylum) are only fully resolved via custom_regex; the built-in segment mode captures the extra-rank content inside the upstream column (§3.5).

Limitation 3. Cycle detection for external taxonomy files excludes adjacent-rank self-loops (e.g., placeholder synonyms such as phylum=SpSt-1190, class=SpSt-1190), but genuine cross-rank cycles still require user correction.

Limitation 4. For trees exceeding 10,000 tips, MRCA computation and nested detection (O(n·k)) can cause substantial delay and high memory use; --low_memory mode alleviates memory pressure via garbage collection, and runtime may grow faster than linearly when the number of candidate groups grows with tree size (tested up to 10,000 tips; the full 189,801-tip bac120 tree remains beyond single-threaded interactive scope). The per-color layer strategy (two ggplot2 layers per group color; 358 layers on the ar53 order-level run) adds noticeable rendering overhead at the order level; this is an inherent constraint of passing colors as non-aesthetic parameters, and single-layer multi-color mapping is listed as future work (§4.4). All benchmarks were obtained on a single machine under the unified protocol of §2; absolute values are hardware-specific.

Limitation 5. The custom Geom objects (GeomPolygonStraight/GeomSegmentStraight) depend on coord$transform() method signatures and ggplot2 internal constants outside the public API, so future ggplot2/ggtree major releases may require adaptation of the rendering layer. Version-compatibility guards warn users at package load time, and vdiffr visual-regression snapshots (tests/testthat/test-straight-vdiffr.R) guard against silent rendering drift across dependency updates; the development environment used ggplot2 4.0.3 and ggtree 4.0.4.

Limitation 6. --multi_tree_mode split currently behaves similarly to all at the output level, distinguishing trees by indexed file names; richer per-tree metadata output is planned (§4.4). Non-monophyletic candidate groups are skipped by default (warn-and-skip; §2.1), so a successful non-strict run may represent a partial collapse. The “displayed leaves after collapse” metric counts visible true tips plus collapsed pseudo-leaf clades and is not directly comparable to tip-count summaries from other tools.

Limitation 7. Geological annotation does not establish that the input tree is a valid chronogram: the branch-length unit must be supplied explicitly (the pipeline aborts otherwise), Rclade warns when root-to-tip distances are highly dispersed, and it never infers divergence times.

### 4.4 Future work

Planned extensions include interactive visualization through plotly/shiny integration, parallelization for very large trees (>50,000 taxa), direct BEAST2 output parsing and node-support handling, additional advanced layouts (e.g., radial), and exploring single-layer multi-color mapping to optimize the per-color layer expansion strategy.

## 5. Software Availability

### Repository

Rclade is publicly available at https://github.com/zengzichao/Rclade (MIT license), with versioned snapshots archived at Zenodo (latest archive snapshot: v1.1.0) (https://doi.org/10.5281/zenodo.22106523 (version 1.1.0; all versions: https://doi.org/10.5281/zenodo.22043060)).

### Installation

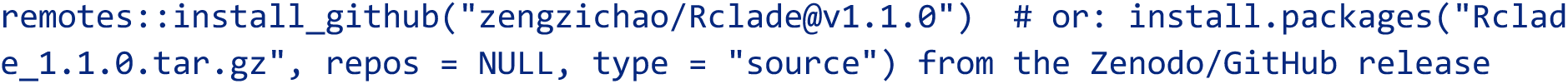

Or use Conda/Docker (development files located in the project root):

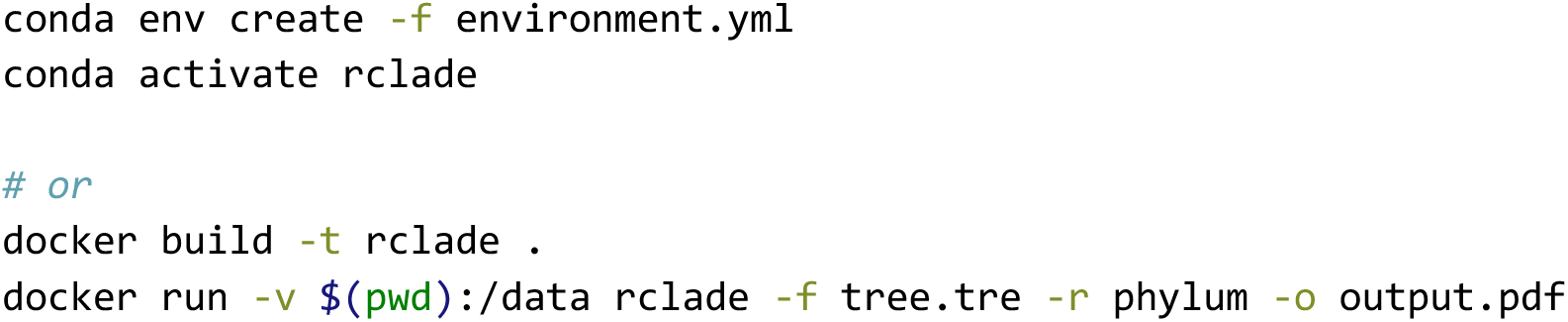

### Version: 1.1.0 (MIT license; requires R ≥ 4.1.0)

#### Documentation

Quick-start guide: vignette("quick_start", package = "Rclade") - Taxonomy- format details: vignette("taxonomy_formats", package = "Rclade") - Publication-ready configuration: vignette("publication_ready", package = "Rclade") - API reference: 149 man pages (26 exported functions accessible via ?function_name)

#### Unit and integration tests

241 test blocks; under CRAN-mode execution 484 expectations pass, 0 fail, and 8 are skipped (including 2 vdiffr visual-regression snapshots that require NOT_CRAN=true; 492 expectations in total), measured with testthat 3.3.1 on R 4.5.3 - Visual-regression snapshots: vdiffr snapshots of collapsed circular/rectangular renderings guard against rendering drift across ggplot2/ggtree updates - Functional workflow tests: covering 17 test categories (see Supporting Information, Functional Test Coverage section) - Self-check: `rclade --check` or `run_rclade_selftest()` - Package check: `R CMD check --as-cran` (R 4.5.3) completes with 0 errors and 0 warnings (only standard packaging-hygiene NOTEs, e.g., CRAN incoming feasibility for a new submission) - CI: GitHub Actions automatically runs R CMD check and coverage checks

#### Reproducibility

All benchmark random seeds are fixed (set.seed(42) in scripts/benchmark_synthetic.R and scripts/benchmark_split_cost.R). The scripts/ directory contains baseline_manual_workflow.R (a complete runnable baseline script demonstrating the equivalent manual workflow; see Supporting Information, Baseline Script section), the benchmark scripts benchmark_synthetic.R, benchmark_split_cost.R, benchmark_real_timing.R (real-data in-session timing, Table 4 source), and run_process_level.sh (unified measurement protocol, §3.2), make_ncbi_rank_shift_example.R (deterministic regeneration of the NCBI rank-shift example), and measure_straightness.R (straight-edge deviation, §3.4); the GTDB real-data parsing-accuracy evaluation script and results (§3.5) are provided in the Zenodo archive. Full sessionInfo() output is provided in benchmark_results/sessionInfo.txt (MacBook Pro, Mac17,2; Apple M5; 32 GB unified memory; macOS 26.6; micromamba r-4.5.3).

#### Example data

 built-in example_tree (50 tips, GTDB-style labels) and polytomy_tree are shipped with the package The GTDB R232 ar53 reference tree (10,122 tips) used in Table 4 and the reformatted 700-tip archaeal timetree of Moody et al. [28] used in Figure 2, Table 4, and §3.5 are available from their original sources (GTDB Release 232 under CC BY-SA 4.0; Moody et al. 2025 under CC BY 4.0). All benchmark result tables remain included in the Zenodo archive (https://doi.org/10.5281/zenodo.22106523 (version 1.1.0; all versions: https://doi.org/10.5281/zenodo.22043060)). The Moody et al. timetree is redistributed with attribution to the original publication (BEAST/Nexus format; tip labels reformatted by us as described in §3.5; topology, branch lengths, and node annotations unchanged); the CC BY 4.0 licence permits further redistribution with attribution to the original publication.

#### CLI wrapper

inst/bin/rclade; add to PATH for direct terminal use:

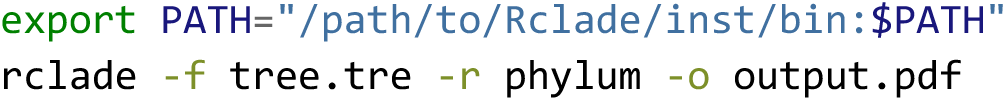

#### Long-term maintenance

the authors commit to maintaining the package for at least 3 years after publication, including bug fixes, dependency updates, and feature extensions.

## Acknowledgements

This work was supported by the National Natural Science Foundation of China (grant nos. 42422209 and 42272354) and the National Key R&D Program of China (grant no. 2023YFC3108600). The authors thank the developers and maintainers of the ggplot2, ggtree, and deeptime packages, on which Rclade directly builds. Computational benchmarks were performed on personal hardware; no high-performance computing resources were used.

## Data Accessibility and Benefit-Sharing

### Data Accessibility Statement

#### Software

The Rclade source code (version 1.1.0, MIT license), user manual (README and three vignettes), and all benchmark and evaluation scripts are archived at Zenodo (https://doi.org/10.5281/zenodo.22106523 (version 1.1.0; all versions: https://doi.org/10.5281/zenodo.22043060)); ongoing development and future releases are maintained at GitHub (https://github.com/zengzichao/Rclade).

#### Benchmark data

All synthetic-benchmark and real-data evaluation results (benchmark_synthetic_rendered.csv, process_level_metrics.csv, benchmark_split_cost_1000.csv, ar53_benchmark.csv, ar53_process_level.txt, straightness_deviation.csv, ncbi_rank_shift_example.csv, taxonomy_accuracy_example_tree.csv) and the full sessionInfo() output are included in the Zenodo archive.

#### Input datasets

The GTDB Release 232 archaeal (ar53; 10,122 tips) and bacterial (bac120; 189,801 tips) reference trees and the ar53 taxonomy table were obtained from the Genome Taxonomy Database (https://gtdb.ecogenomic.org/downloads; Parks et al. [1]; used under the Creative Commons Attribution-ShareAlike 4.0 International (CC BY-SA 4.0) licence, and the derived tree and taxonomy files are therefore available from the original source under CC BY-SA 4.0). The 700-tip archaeal timetree is that of Moody et al. [28] (tip labels reformatted by us; topology, branch lengths, and node annotations unchanged; available from the original publication under the Creative Commons Attribution 4.0 International (CC BY 4.0) licence; the exact reformatted file used in this study is archived with the package (inst/extdata/moody2025_gbm_laca_timetree.nwk, SHA-256 in MANIFEST.tsv)).

#### Benefit-Sharing Statement

This computational study did not collect new biological material or access restricted genetic resources. It used publicly available phylogenetic and taxonomic datasets under their stated licenses; provenance and attribution are reported in the Data Accessibility Statement. Benefits are shared through the open-source software, reproducible workflows, and public benchmark materials.

#### Author Contributions

Zichao Zeng: Conceptualization, Methodology, Software, Validation, Investigation, Data curation, Visualization, Writing – original draft. Yinzhao Wang: Supervision, Funding acquisition, Writing – review & editing. All authors have read and approved the final manuscript.

