## Supporting Information for "Rclade: automated taxonomic collapsing and geological-timescale annotation of time-calibrated phylogenetic trees in R"

**Table S1. Key parameters of `plot_timetree()`** (complete reference: `?plot_timetree`; 149 man pages cover all exported functions).

| Parameter | Type | Default | Description |
|---|---|---|---|
| `tree` | phylo / multiPhylo / character | — (required) | Input tree object or file path (Newick/Nexus/BEAST auto-detected) |
| `tree_index` | integer | NULL | Tree index for multi-tree files |
| `multi_tree_mode` | character | "error" | Multi-tree handling: error / first / all / split |
| `rank` | character | "none" | Collapsing rank (phylum, class, order, …) or "none" |
| `groups` | list | NULL | Manual group definitions (named tip lists; overrides rank parsing) |
| `layout` | character | "rectangular" | Layout: rectangular / circular |
| `triangle_mode` | character | "mixed" | Triangle mode for collapsing (forced to "max" in circular layout) |
| `space_mode` | character | "proportional" | Clade space allocation: proportional / equal |
| `taxonomy_format` | character | "auto" | GTDB / Silva / NCBI / embedded / custom_rank / custom_regex / auto |
| `custom_patterns` | list | NULL | Per-rank regex patterns (required for custom_regex) |
| `taxonomy_delimiter_mode` | character | "reverse" | Embedded parsing strategy: reverse / greedy / segment |
| `taxonomy_file` | character | NULL | External taxonomy file path (label → taxonomy) |
| `taxonomy_file_priority` | logical | TRUE | File taxonomy overrides label-based parsing |
| `color_palette` | character | "viridis" | Palette name (viridis / plasma / Set1 / custom, …) |
| `color_mapping` | named character | NULL | Explicit group → color mapping |
| `color_rank` | character | NULL | Rank used for branch coloring |
| `add_timescale` | logical | TRUE | Add geological timescale |
| `timescale_mode` | character | "radial" | Timescale mode: radial / linear |
| `timescale_levels` | character | c("eras", "eons") | Timescale strip datasets |
| `timescale_version` | character | "ICS 2023/02" | International Chronostratigraphic Chart version |
| `unit` | character | NULL | Time unit (Ma / Ga); NULL leaves native units untouched and is only valid with add_timescale = FALSE; when add_timescale = TRUE the pipeline aborts unless unit is supplied explicitly (Rclade does not infer units) |
| `geo_events` | logical | FALSE | Annotate geological events (e.g., GOE/NOE) |
| `show_tip_labels` | logical | FALSE | Show tip labels |
| `tip_label_size` | numeric | 2 | Tip label font size |
| `line_width` | numeric | 1 | Branch line width |
| `legend_position` | character | "bottom" | Legend position (bottom / right / none / …) |
| `legend_nrow` / `legend_ncol` | integer | NULL | Legend layout control |
| `show_clade_label` | logical | FALSE | Label collapsed clades |
| `show_clade_count` | logical | TRUE | Show tip counts in clade labels |
| `clade_label_offset` | numeric | 50 | Clade label offset |
| `show_support` | logical | FALSE | Show node support values |
| `show_hpd` | logical | FALSE | Show 95% HPD intervals (BEAST trees) |
| `strict` | logical | FALSE | Escalate non-monophyly warnings to errors |
| `highlight` | list | NULL | Special-node highlighting (e.g., LUCA/LACA/LBCA) |
| `low_memory` | logical | FALSE | Trigger garbage collection for large trees |
| `ignore_malformed` | logical | FALSE | Tolerate malformed labels |
| `output` | character | NULL | Output path (pdf/png/svg/tiff/eps inferred from extension) |
| `overwrite` | character | "ask" | Overwrite policy for existing outputs |
| `width` / `height` | numeric | 14 / 10 | Output figure size (inches) |
| `opts` | rclade_options / list | NULL | Options object supplying defaults (CLI/config integration) |

Note that monophyly checking is always enabled inside the pipeline (warn-and-skip by default: a non-monophyletic group produces a warning and is omitted from the collapse plan; escalate via strict = TRUE) and is therefore not a separate user-facing switch. Per-group status (total/collapsed/singleton/skipped) is reported in rclade_info and the completion log.

#### Supplementary Figures

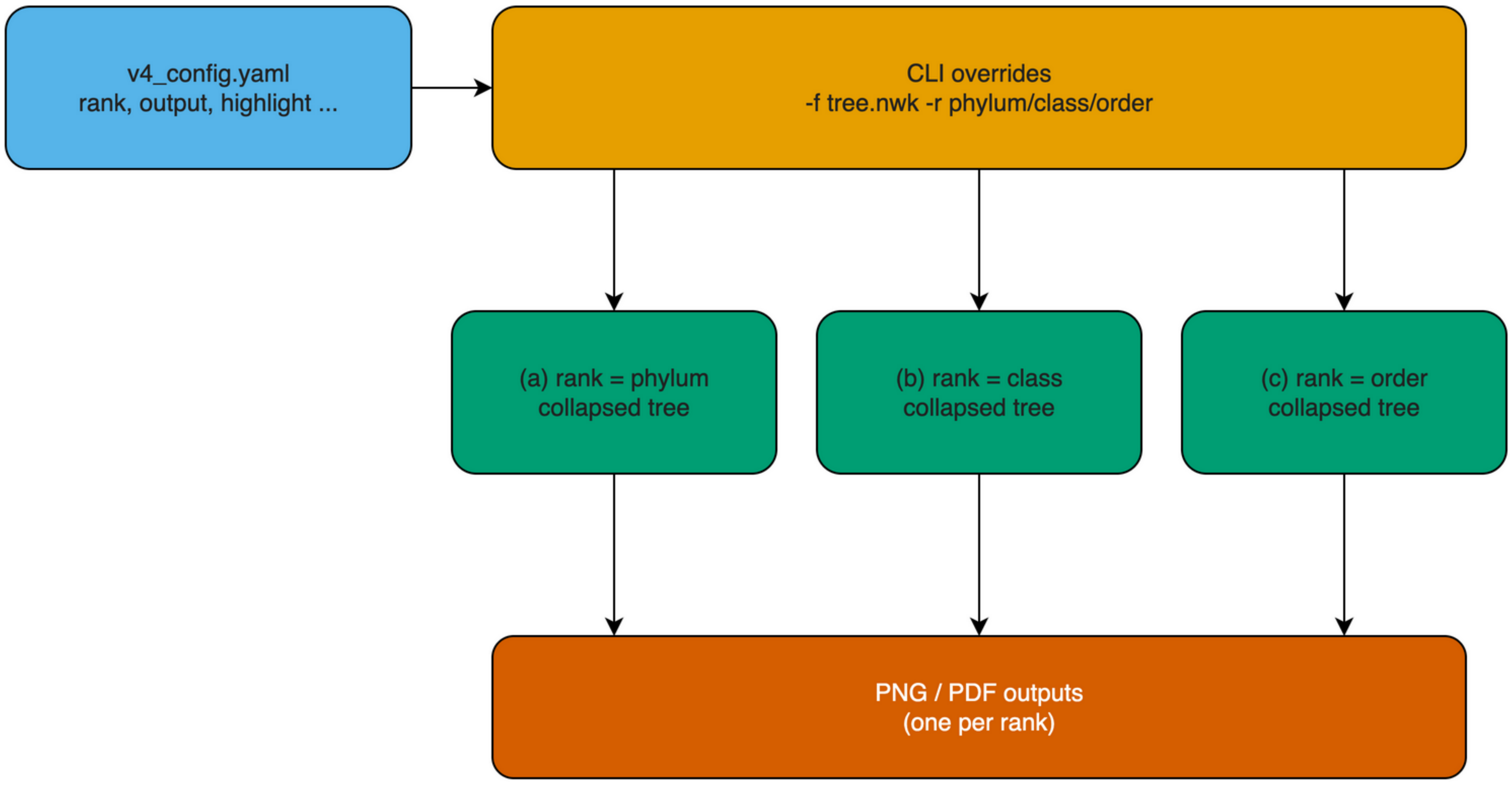


**Figure S1. YAML configuration-driven workflow.** Using the same `v4_config.yaml` as the default and overriding only the `-r` parameter via the CLI to generate phylum-, class-, and order-level collapsing results.

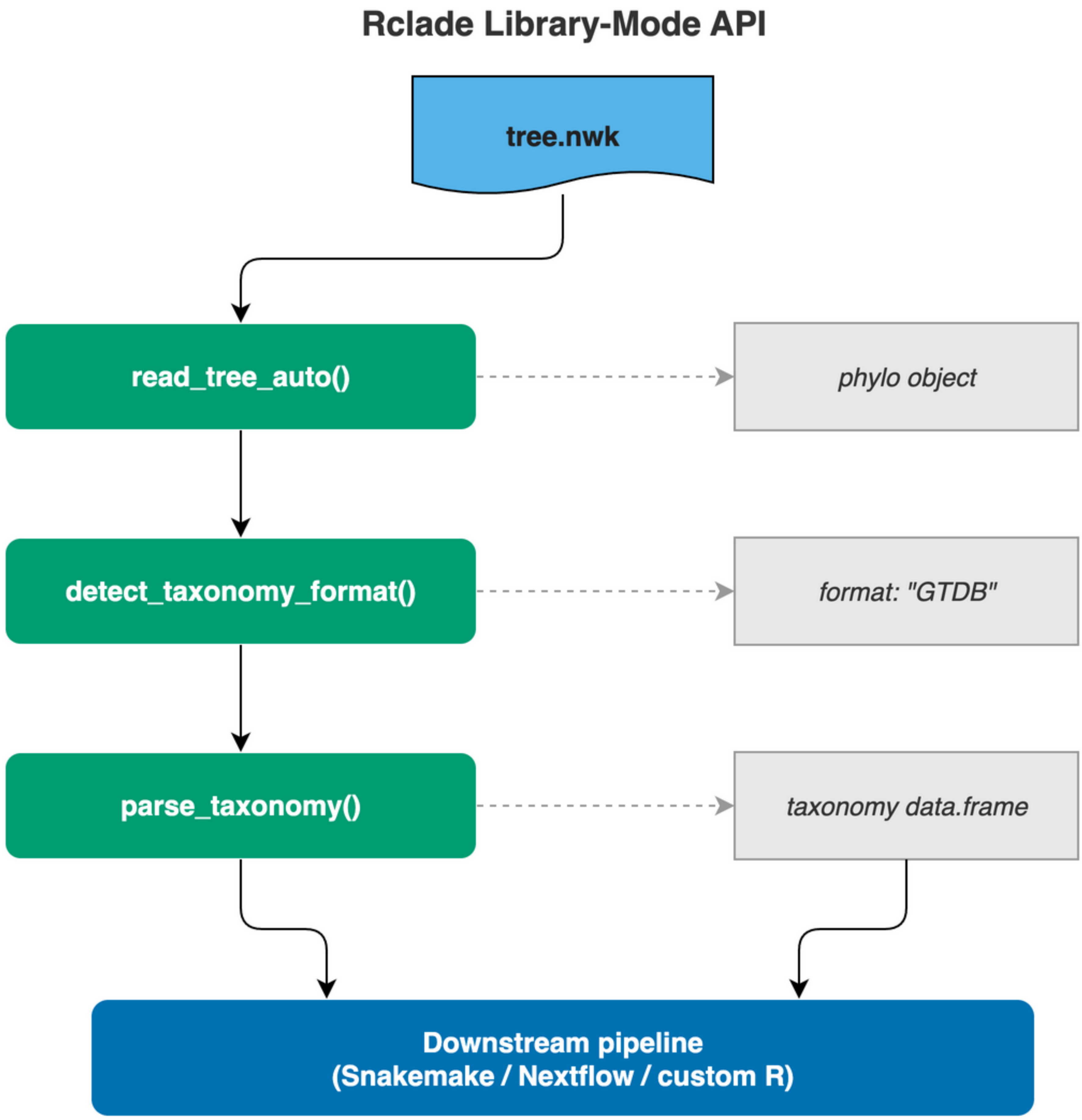


**Figure S2. Library-mode API example.** Showing the combined use of `read_tree_auto()`, `detect_taxonomy_format()`, and `parse_taxonomy()`; parsed results can feed directly into downstream pipelines.

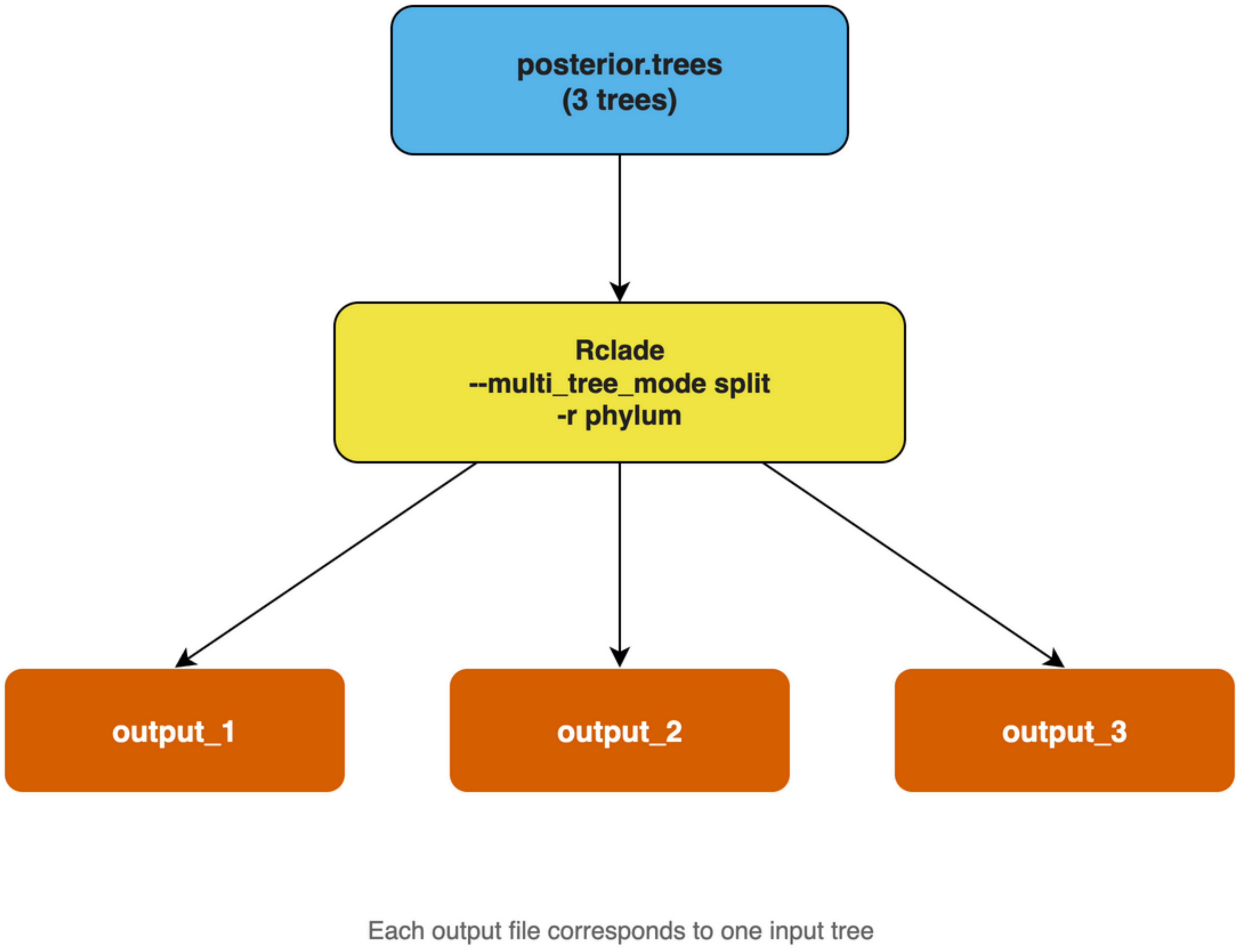


**Figure S3. Multi-tree split mode.** An input posterior file containing 3 trees is processed with `--multi_tree_mode split`, producing one independent output file per tree for downstream parallel dispatching.

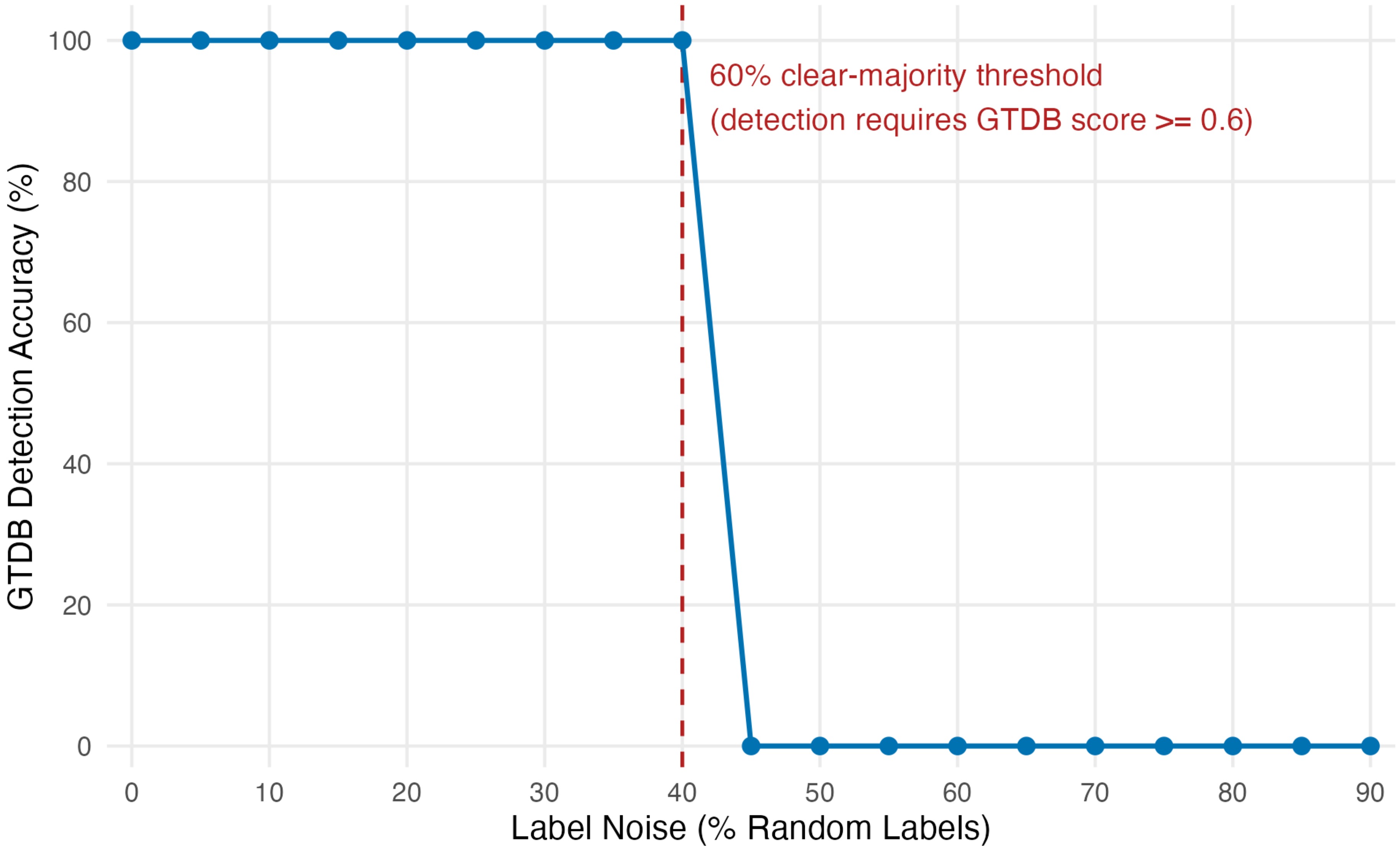


**Figure S4. Deterministic sensitivity of the GTDB decision rule to idealized label replacement (illustrative; not an empirical accuracy estimate).** Synthetic GTDB-style labels were mixed with random labels (0%–90%; the noise experiment was conducted on GTDB-style labels), and the detection outcome was recorded using Rclade::detect_taxonomy_format(). The red dashed line marks the 60% clear-majority threshold: a format is accepted only when its prefix-match score is strictly above 0.5 and at least 0.1 away from the tie line (score ≥ 0.6); with noise fraction f the GTDB score equals 1 − f, so detection succeeds while noise ≤ 40% and falls back to “unknown” beyond.

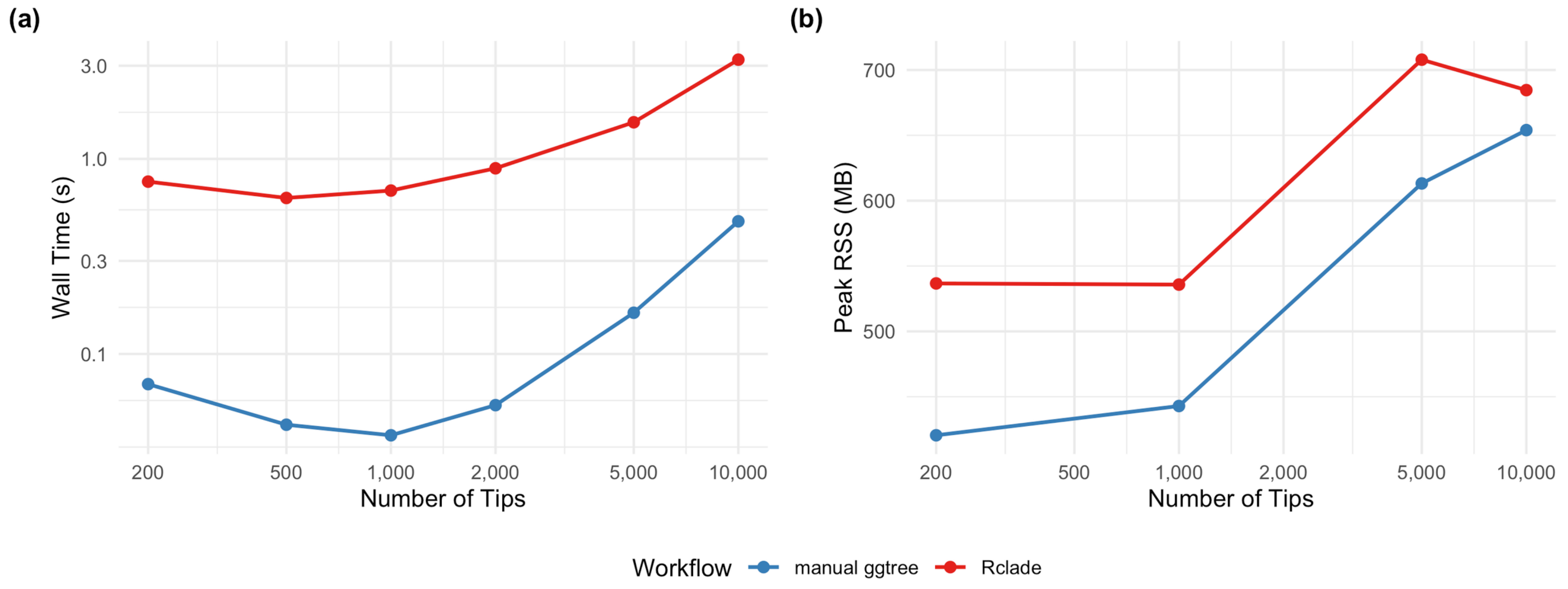


**Figure S5. Performance scaling curves on synthetic datasets.** Left: rendering time (log scale) for Rclade versus a manual ggtree baseline; right: peak RSS memory. These curves report in-session rendering medians from `bench::mark` (warm R session, excluding process startup and package loading; 5 replicates at every scale, unified 2026-08 protocol); Table 3 of the main text reports both this in-session series and the process-level wall-clock measured with `/usr/bin/time`, which is systematically larger for the same workloads because it includes R startup and package loading (see the measurement-level note below Table 3).

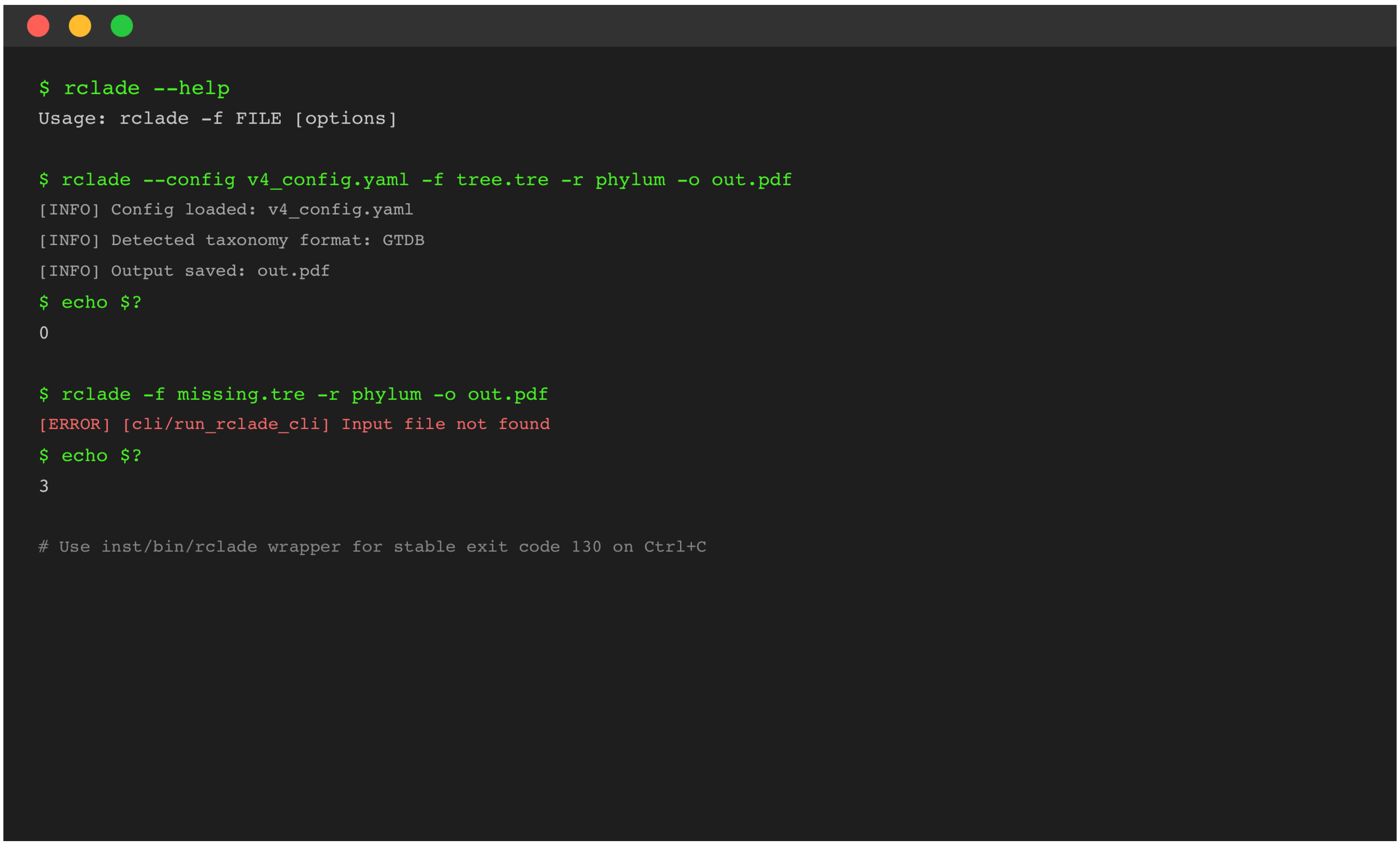


**Figure S6. CLI, configuration, and exit-code example.** Terminal-style illustration showing `rclade --help`, config-file usage, success/input-error exit codes, and the `inst/bin/rclade` wrapper's stable SIGINT exit-code 130 support.

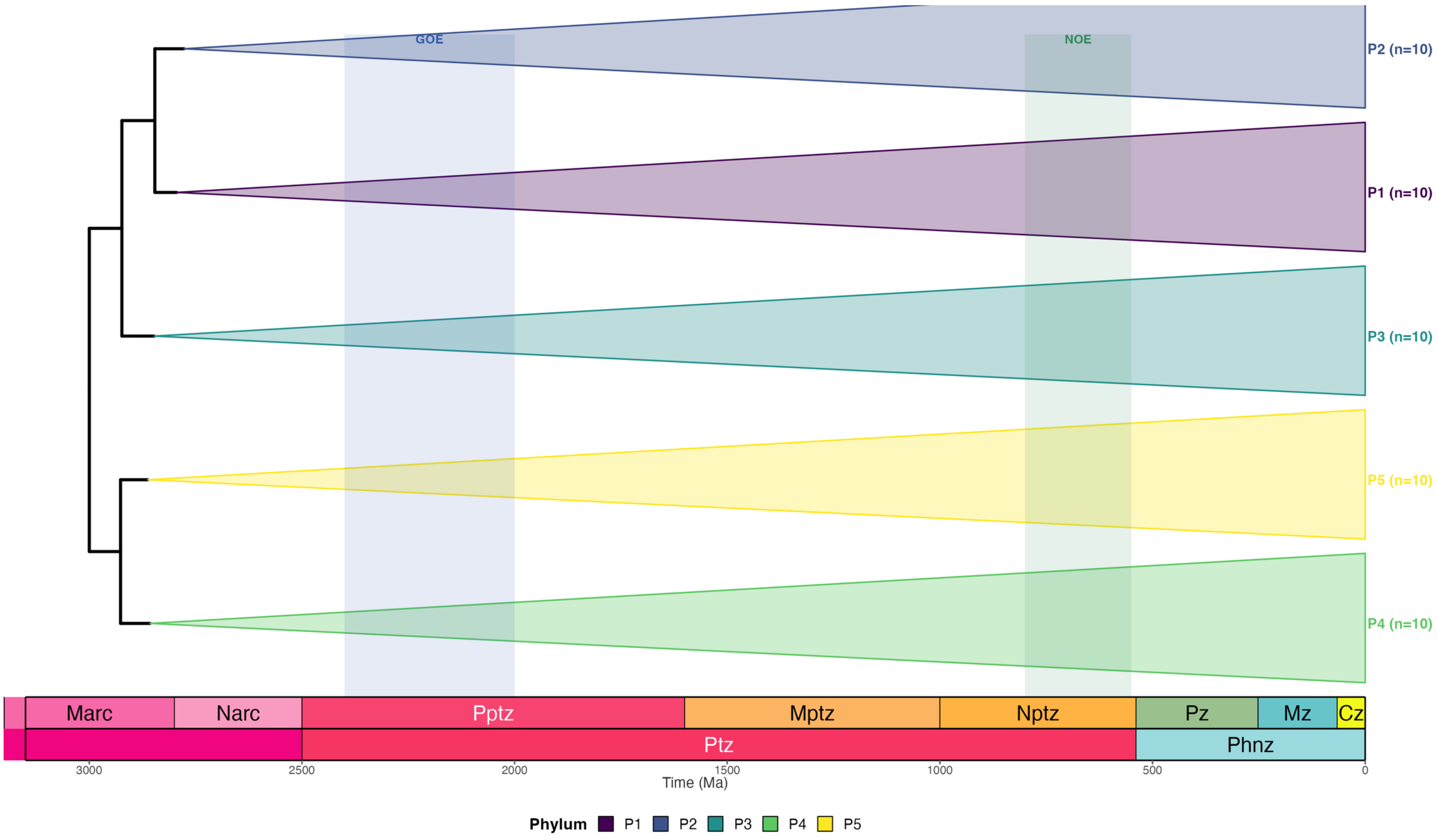


**Figure S7. Built-in example_tree phylum-level collapsing example.** Phylum-level taxonomic collapsing of Rclade's built-in `example_tree` dataset, demonstrating the combined use of taxonomic label parsing, geological timescale integration, geological event annotation (GOE/NOE), and clade labelling.

#### Supplementary Methods: Production-Grade Engineering Practices

##### Temporary-file permissions

In shared HPC environments, temporary files may contain sensitive data. Rclade's `managed_tempfile()` sets permissions to `0600` immediately after creation, and `managed_tempdir()` sets permissions to `0700`, best-effort preventing other users from reading them (the chmod follows file creation; behavior is platform- and filesystem-dependent).

##### CI coverage threshold

`codecov.yml` sets overall and patch coverage targets to 85% and marks core modules (parse-taxonomy, taxonomy-file, validate-deep, monophyly, compute-mrca, read-input) individually. The CI workflow keeps fail_ci_if_error: false (no CODECOV_TOKEN configured); coverage is therefore a monitoring metric, not an enforced quality gate.

##### Structured logging

The logger supports a `.module` argument and outputs `[MODULE/FUNCTION]` tags (e.g., `[validate-deep/check_name_safety]`, `[compute-mrca/compute_mrca_map]`) so pipeline logs can quickly locate problems.

##### Standard exit codes and wrapper script

The Rclade CLI uses an application-level exit-code contract: 0 (success), 1 (runtime error), 2 (parameter error), 3 (input-data error), 130 (user interrupt/SIGINT). Codes 0 and 130 follow common Unix conventions; codes 2 and 3 are Rclade-specific categories. The inst/bin/rclade shell wrapper captures SIGINT and correctly propagates exit code 130.

##### Strip Annotations

The `--strip_annotations` CLI option removes node annotation information (bootstrap support values, NHX metadata) from tree files before rendering, which may reduce output file sizes (magnitude not quantified).

##### Logger concurrency limitation

Rclade's logging system uses a package-level global environment and does not support concurrent multi-process writes to the same log file. In parallel scenarios, users should assign independent log files via `--log_file` (e.g., `--log_file logs/task_$SLURM_JOB_ID.log`).

##### Searched alternatives

To substantiate the claim in main text §2.7 that "to our knowledge, no built-in parameter or existing extension addresses this rendering artifact as of ggtree 4.0.4", we systematically searched the following sources for potential alternatives (search time-point: ggtree 4.0.4 / deeptime 2.4.0, verified 2026-08):

**Table S2. Searched alternatives and reasons for exclusion.**

| Candidate | Source | Reason it does not solve the problem |
|---|---|---|
| `ggtree::collapse()` parameters | ggtree GitHub issue tracker / documentation | A targeted search of the ggtree documentation and the YuLab-SMU/ggtree issue tracker found no parameter to disable `coord_munch()` interpolation and no closed issue providing a fix for curved collapsed-triangle edges under polar coordinates as of v4.0.4 |
| `ggplot2::annotate()` / `ggtree::geom_hilight()` | ggplot2/ggtree documentation | Uses annotation layers for highlighting, not collapsing semantics; does not reduce tip count or produce collapsed triangles; cannot substitute for `collapse()` functionality |
| `ggforce::geom_shape()` / `geom_mark_*()` | ggforce package documentation | General-purpose shape drawing; no phylogenetic collapsing semantics; does not address the `coord_munch()`-induced straight-edge problem under polar coordinates |
| `ggforce::geom_shape()` + hand-computed polar vertices | ggforce + manual vertex computation | Can draw straight polygons under `coord_polar()` if triangle vertices are pre-computed by hand, but requires reimplementing MRCA geometry, tip-spacing vertex correction, panel-clipping control, and color-scale conflict handling for every plot; no reusable collapsing semantics, no automation |
| `ggtreeExtra::geom_fruit()` | ggtreeExtra documentation | Overlays data annotation layers outside the tree ring; not intra-branch collapsing; does not alter tree topology display or reduce visual clutter |
| Manual vertex interpolation workarounds | StackOverflow / `coord_munch` discussions | Only scattered manual vertex-interpolation workarounds exist (e.g., hand-computing polar vertices); no reusable package/extension; does not handle panel clipping, color-scale conflicts, or other engineering concerns |

In summary, as of the search time-point, no built-in parameter or reusable extension in the R ecosystem simultaneously addresses straight-edge rendering of polar-coordinate collapsed triangles, panel-clipping integrity, and color-scale conflict. Rclade's custom ggproto approach fills this gap.

### Straight-Edge Rendering of Collapsed Triangles in Circular Layout

#### 1. Problem Background

In circular (fan) layouts of phylogenetic trees, when `ggtree::collapse()` is used to collapse monophyletic clades into triangles, the triangle edges appear as curves rather than straight lines. This occurs because `ggtree` internally uses `geom_polygon()` to draw the collapsed triangles, and `geom_polygon()` delegates to `GeomPath`, whose `draw_panel()` method calls `coord_munch()` to interpolate straight line segments into curves under polar/radial coordinate systems.

**Effect**: The two slanted edges of the collapsed triangle (connecting the MRCA node to the outer vertices) become curved arcs, producing a "flame-like" or "wavy" appearance that severely degrades visual quality and readability.

**This issue remains unresolved in `ggtree` (v4.0.4)**. `ggtree::collapse()` provides no parameter to disable curve interpolation in polar coordinates.

#### 2. Root Cause Analysis

##### 2.1 How `coord_munch()` Works

`ggplot2`'s `coord_munch()` is an internal function for polar coordinate systems. Its purpose is to interpolate a straight line segment in data space into many small segments, simulating the "correct" curve under polar coordinates.

```
Data space (Cartesian)     Polar transform     Render space (Cartesian npc)
  V1 ──────────── V2   →   coord_munch()   →   V1 ⌒⌒⌒⌒ V2
  (straight)               (interpolated)      (arc)
```

For `geom_polygon()`, the `GeomPath$draw_panel()` workflow is: 1. Extract polygon vertex coordinates. 2. Call `coord_munch()` to insert intermediate points between each pair of edge endpoints. 3. Call `coord$transform()` for each intermediate point to convert to npc space. 4. Connect all intermediate points with `polylineGrob`.

Under polar coordinates, intermediate points of a straight segment are mapped to different angles and radii, ultimately forming an arc in npc space.

##### 2.2 Why Rectangular Layout Is Not Affected

In rectangular layouts, the coordinate system is standard Cartesian, and `coord_munch()` does not bend straight lines (because the transformation is linear). Therefore, `geom_polygon()` draws normal straight-edged triangles.

#### 3. Solution

##### 3.1 Core Idea: Bypass `coord_munch()`

We created two custom `ggproto` Geom objects—`GeomPolygonStraight` and `GeomSegmentStraight`—that call `coord$transform()` **only on vertices/endpoints**, and then use `grid` primitives (`polygonGrob` / `segmentsGrob`) to draw **straight-edged** glyphs in npc space.

```
Data space (Cartesian)     Direct transform    Render space (Cartesian npc)
  V1 ──────────── V2   →   transform(V1)   →   V1 ──────── V2
  (straight)               transform(V2)       (straight!)
                           (no interpolation)
```

Because `grid::polygonGrob()` and `grid::segmentsGrob()` draw true straight lines in npc space without further coordinate transformation, the final rendering shows straight edges.

##### 3.2 `GeomSegmentStraight` Implementation

```r
GeomSegmentStraight <- ggplot2::ggproto(
  "GeomSegmentStraight", ggplot2::Geom,
  required_aes = c("x", "y", "xend", "yend"),
  draw_panel = function(data, panel_params, coord, ...,
                        segment_colour = "black", na.rm = FALSE) {
    # Split start and end points
    starts <- data; ends <- data
    ends$x <- data$xend; ends$y <- data$yend
```

```r
    # Transform endpoints only — do NOT call coord_munch()
    starts_t <- coord$transform(starts, panel_params)
    ends_t   <- coord$transform(ends,   panel_params)

    # Draw straight segments in npc space
    grid::segmentsGrob(
      x0 = starts_t$x, y0 = starts_t$y,
      x1 = ends_t$x,   y1 = ends_t$y,
      gp = grid::gpar(col = segment_colour, ...)
    )
  }
)
```

**Key points**: - `coord$transform()` transforms only endpoints, without interpolating intermediate points. - `segmentsGrob()` draws a straight line between two points in npc space. - Color is passed via the non-aesthetic parameter `segment_colour` to avoid conflict with `scale_color_manual`.

##### 3.3 `GeomPolygonStraight` Implementation

```r
GeomPolygonStraight <- ggplot2::ggproto(
  "GeomPolygonStraight", ggplot2::Geom,
  required_aes = c("x", "y"),
  draw_panel = function(data, panel_params, coord, ...,
                        polygon_fill = "grey50", polygon_alpha = 0.3,
                        na.rm = FALSE) {
    # Transform vertices only — do NOT call coord_munch()
    data_t <- coord$transform(data, panel_params)

    # One polygonGrob per group (each group = one triangle)
    grobs <- lapply(split(data_t, data_t$group), function(sub) {
      grid::polygonGrob(
        x = sub$x, y = sub$y,
        gp = grid::gpar(fill = polygon_fill, alpha = polygon_alpha, col = NA)
      )
    })
    do.call(grid::gList, grobs)
  }
)
```

**Key points**: - `polygonGrob()` connects vertices with straight lines in npc space, forming straight-edged polygons. - `col = NA` means no border is drawn; borders are rendered separately by `GeomSegmentStraight`. - Fill color is passed via the non-aesthetic parameter `polygon_fill`.

##### 3.4 Color Conflict and Resolution

`ggtree::collapse()` passes fill and border colors as aesthetic parameters (`aes_params`) to its internal `geom_polygon()`. When `scale_fill_manual` is also used to control the legend, fixed color values conflict with the discrete color scale, causing a "`No shared levels found`" warning and a missing legend.

**Solution**: Call `ggtree::collapse()` with `fill = NA, color = NA`, making its internal `geom_polygon` completely invisible (used only for data modification—collapsing descendant nodes and reassigning y coordinates). Then overlay visible fill and border layers with `GeomPolygonStraight` and `GeomSegmentStraight`, passing colors via non-aesthetic parameters to fully bypass the ggplot2 scale system.

```r
# ggtree::collapse() performs only data modification, no visible drawing
ggtree::collapse(p, node = node_id, mode = "max",
                 fill = NA, color = NA)

# Then overlay straight-edged triangles with custom Geoms
p <- p + geom_polygon_straight(data = verts, aes(x, y, group = group),
```

```
                                polygon_fill = group_color, polygon_alpha = 0.3)
p <- p + geom_segment_straight(data = segs, aes(x, y, xend, yend),
                                segment_colour = group_color)
```

#### 4. Triangle Geometry Calculation

##### 4.1 When to Compute Vertices

Triangle vertices **must be computed before calling `ggtree::collapse()`**, because `collapse()` reassigns the y coordinates of descendant nodes in `p$data`, changing their positions.

##### 4.2 Vertex Position Correction

`ggtree::get_clade_position()` returns `ymin`/`ymax` using a fixed 0.5 offset. In circular layouts, `scaleClade()` compresses the y range, so a fixed 0.5 offset is too large relative to the compressed tip spacing, causing vertices to fall outside the coordinate scale range and be censored to NA.

**Correction**: Replace the fixed 0.5 offset with half of the actual (post-`scaleClade`) tip spacing:

```
tip_spacing <- median(diff(sort(unique(tip_y))))
offset <- tip_spacing * 0.5
ymin <- sp$ymin + 0.5 - offset  # = min(clade_tip_y) - offset
ymax <- sp$ymax - 0.5 + offset  # = max(clade_tip_y) + offset
```

##### *4.3 Forcing "max" Mode in Circular Layout*

`ggtree::collapse()` supports three triangle modes, but only `"max"` produces the correct visual effect under polar coordinates:

**Table S3. Triangle modes in ggtree::collapse() and their polar effects.**

| Mode | Vertex positions | Polar effect |
|---|---|---|
| `max` | V1=(x_node, y_node), V2=(xmax, ymin), V3=(xmax, ymax) | Correct inward-pointing wedge (MRCA vertex near center, wide edge at outer rim) |
| `min` | V1=(x_node, y_node), V2=(xmin, ymin), V3=(xmin, ymax) | Outward-pointing wedge (MRCA vertex at outer rim, wide edge near center) |
| `mixed` | V1=(x_node, y_node), V2=(xmin, ymin), V3=(xmax, ymax) | Diagonal slash crossing adjacent clade territory |

Therefore, in circular layouts, regardless of the user-specified `triangle_mode`, the geometric computation forces `"max"` mode:

```
eff_mode <- if (layout == "circular" && mode != "none") "max" else mode
```

##### 4.4 Rendering integrity: panel clipping control

Even after the correction in Section 4.2 of this Supplement keeps triangle vertices within the coordinate scale range, and after Sections 4.1–4.3 of this Supplement produce straight-edge geometry, the collapsed triangle's **MRCA vertex (V1) may still lie outside the original y-axis panel bounding box computed from tip nodes** (e.g., panel y range is `[0, 47.125]`, but the root clade's triangle vertex y can reach `50`). ggplot2's default `clip = "on"` clips content outside the panel, causing the **top triangle apex to be truncated**.

**Fix**: Set `coord_cartesian(clip = "off")` in the coordinate system returned by the theme layer (`theme_timetree()`), disabling panel clipping so that collapsed geometry extending beyond the panel boundary is rendered completely:

```
# theme_timetree() returns list(theme, coord), where
coord <- ggplot2::coord_cartesian(clip = "off")
# In pt_step7_finalize_plot(): p <- p + theme_result$theme + theme_result$coord
```

**Note**: The return value of `theme_timetree()` changed from a single ggplot2 theme object to `list(theme, coord)` as of this fix; direct callers should use `p + th$theme + th$coord`; `plot_timetree()` has been adapted internally, so ordinary users need not change anything. This is a breaking change, recorded in `NEWS`.

This fix complements the adaptive vertex-position correction in Section 4.2 of this Supplement: the former addresses vertices being censored to `NA` by the oob handler, while the latter addresses vertices being clipped outside the panel bounding box. Together they ensure **complete, undistorted rendering** of collapsed triangles.

#### 5. Complete Workflow

Execution flow of `collapse_by_groups()`:

```
Step 1: scaleClade (all clades, depth-first)
  ├── Compress y range of each clade (proportional or equal mode)
  └── Done before any collapse to avoid subsequent clade position changes

Step 2: Collapse clades one by one (depth-first)
  for each clade (deepest first):
    ├── compute_collapse_vertices()  ← compute vertices BEFORE collapse
    ├── compute_collapse_segments()  ← compute edge segment endpoints
    ├── ggtree::collapse(fill=NA, color=NA)  ← data modification only
    └── collect vertex/segment data into polygon_list / segment_list

Step 3: Overlay straight-edged glyphs (circular layout)
  ├── One geom_polygon_straight layer per color
  └── One geom_segment_straight layer per color
```

##### *5.1 Depth-First Ordering*

Collapsing must start from the deepest clade (`sort_by_depth()`), because after a clade is collapsed, its descendant nodes are repositioned in `p$data` and can no longer be accessed via original node IDs. Depth-first ordering ensures internal clades are collapsed before external ones.

##### *5.2 One Layer per Color*

ggplot2's scale system does not support multi-color mapping within a single layer when color is passed as a non-aesthetic parameter. Therefore, a separate `geom_polygon_straight` / `geom_segment_straight` layer is created for each color:

```r
for (uc in unique(all_polys$fill_colour)) {
  poly_subset <- all_polys[all_polys$fill_colour == uc, ]
  p <- p + geom_polygon_straight(data = poly_subset,
                                aes(x, y, group = group),
                                polygon_fill = uc, polygon_alpha = 0.3)
}
```

#### 6. Code Location

All implementations are in R/collapse-workflow.R of the package source (package version 1.1.0, archived at Zenodo, https://doi.org/10.5281/zenodo.22106523 (version 1.1.0; all versions: https://doi.org/10.5281/zenodo.22043060); line numbers intentionally omitted because they drift across releases):

**Table S4. Package components and their descriptions.**

| Component | Description |
|---|---|
| `GeomSegmentStraight` | Straight-segment Geom (bypasses coord_munch) |
| `GeomPolygonStraight` | Straight-edged polygon Geom (bypasses coord_munch) |
| `compute_collapse_vertices()` | Triangle vertex computation (includes tip spacing correction) |
| `compute_collapse_segments()` | Triangle edge segment endpoint computation |
| `collapse_by_groups()` | Main function: scaleClade → collapse → overlay straight glyphs |

A quantitative validation of the straight-edge effect (maximum perpendicular deviation of triangle edges from the ideal chord in npc space, measured on the Figure 3 tree) is provided by `scripts/measure_straightness.R` (main text §3.4; benchmark_results/straightness_deviation.csv).

#### 7. Effect Comparison

**Table S5. Rendering effect comparison before and after the fix.**

| Before Fix | After Fix |
|---|---|
| Triangle edges are arcs (flame/wave shape) | Triangle edges are straight |
| Fill color conflicts with `scale_fill_manual` | Colors passed via non-aesthetic parameters, no conflict |
| Vertices may be censored to NA by oob handler | Vertex positions adapt to tip spacing |
| Top triangle apex truncated by panel clipping | Panel clipping disabled (clip = "off"), vertices fully visible (§4.4) |

| Before Fix | After Fix |
|---|---|
| mixed/min modes produce incorrect geometry | Circular layout forces max mode |

##### 8. Limitations

1. **Applies only to triangles from `ggtree::collapse()`**: Other `ggtree` functions (e.g., `geom_motif`, `geom_cladelab`) would require similar treatment if they exhibit curve issues under polar coordinates.
2. **One layer per color**: When many collapsed clades have distinct colors (e.g., 100+ phyla), a large number of layers is created, which may affect rendering performance.
3. **Generality of `coord_munch()` bypass**: This method can theoretically be applied to any ggplot2 Geom that needs straight lines under polar coordinates, but a corresponding `ggproto` variant must be written for each Geom type.
4. **Non-aesthetic parameter pass-through**: Colors are passed to the custom Geoms as non-aesthetic parameters (`polygon_fill`, `segment_colour`); user code that additionally injects color-related arguments through `...` may conflict with these parameters. This is the price of bypassing the ggplot2 scale system, and is why per-color layers are used instead of aesthetic mapping.

#### CLI Interface Reference

The Rclade command-line interface is invoked via `Rscript -e "Rclade::run_rclade_cli()"` or through the `inst/bin/rclade` shell wrapper. The interface accepts the following primary arguments: `-f` / `--file` (input tree file path, required), `-r` / `--rank` (taxonomic rank for collapsing, e.g., phylum, class, order), `-o` / `--output` (output file path), `-l` / `--layout` (rectangular or circular), `--config` (YAML configuration file), `--taxonomy_format` (GTDB, Silva, NCBI, embedded, custom_rank, custom_regex, or auto), `--timescale_mode` (radial, linear), `--color_palette` (viridis, plasma, Set1, etc.), `--multi_tree_mode` (first, all, split), `--strict` (escalate monophyly warnings to errors), `--low_memory` (trigger garbage collection for large trees), `--strip_annotations` (remove node annotations before rendering), `--log_level` (DEBUG, INFO, WARNING, ERROR), and `--log_file` (redirect log output to file). The `--check` flag runs a built-in self-test suite without requiring an input tree. Exit codes follow the Rclade application-level contract (codes 0 and 130 follow common Unix conventions; 2 and 3 are Rclade-specific): 0 (success), 1 (runtime error), 2 (parameter error), 3 (input-data error), and 130 (user interrupt via SIGINT).

#### Functional Test Coverage

Rclade's test suite covers 17 functional categories spanning the complete pipeline: tree reading and format detection (Newick, Nexus, BEAST XML, multi-tree), deep input validation (malformed Newick, control characters, BiDi markers, empty files, duplicate tips, negative branch lengths, self-loops), taxonomic-format detection and parsing (all five formats with noise injection), MRCA computation and nested-conflict resolution, monophyly checking (strict and permissive modes), geological-timescale integration (unit switching, event annotation), color-palette generation and legend layout, batch collapsing with depth-first ordering, straight-edge rendering geometry (vertex computation, clip-off behavior), configuration-file loading and parameter priority, library-mode API functions (read_tree_auto, parse_taxonomy, detect_taxonomy_format), multi-tree split mode, CLI argument parsing and exit codes, interrupt handling and resource cleanup, output file saving (PDF, PNG, SVG with overwrite protection), external taxonomy-file merging with cycle detection, and encoding normalization (UTF-8, BOM, line endings). The test suite comprises 241 test blocks; under CRAN-mode execution, 484 expectations pass with 0 failures and 8 skips (including 2 vdiffr snapshot tests that require NOT_CRAN=true; 492 expectations in total), measured with testthat 3.3.1 on R 4.5.3. The vdiffr visual-regression snapshots (tests/testthat/test-straight-vdiffr.R) guard the collapsed-triangle rendering output against dependency drift.

#### Test Environment

All benchmarks and tests were conducted on a MacBook Pro (Mac17,2) with an Apple M5 chip and 32 GB unified memory, running macOS 26.6 (Tahoe). The R environment was managed via micromamba (r-4.5.3). Key package versions: R 4.5.3, ape 5.8-1, ggtree 4.0.4, ggplot2 4.0.3, deeptime 2.4.0, rlang 1.2.0, stringr 1.6.0, tidytree 0.4.8, viridisLite 0.4.3, phangorn 2.12.1, treeio 1.34.0, bench 1.1.4, vdiffr 1.0.9, testthat 3.3.1. The complete `sessionInfo()` output is provided in `benchmark_results/sessionInfo.txt` of the package repository.

#### Baseline Manual Workflow Script

The following script demonstrates the traditional multi-package approach to GTDB-format tree collapsing with geological timescale, an illustrative manual workflow (not functionally equivalent to Rclade's single plot_timetree() call: it omits format detection, strict/skip policies, nested ordering, and legend/timescale semantics) (~100 lines coordinating five packages). The core-collapsing-only baseline used for the overhead assessment in Table 3 of the main text is a stripped-down version of steps 3–6 (MRCA computation, scaleClade, collapse, rendering), embedded in scripts/benchmark_synthetic.R.

```r
library(ape)
library(ggtree)
library(ggplot2)
library(deeptime)
library(viridisLite)

# 1. Read tree
tree <- read.tree("tree.nwk")
labels <- tree$tip.label

# 2. Parse GTDB taxonomy (extract phylum)
parse_gtdb_phylum <- function(lab) {
  m <- regmatches(lab, regexec("p__([^;]+)", lab))[[1]]
  if (length(m) >= 2) m[2] else NA_character_
}
phylum_vec <- vapply(labels, parse_gtdb_phylum, character(1))

# 3. Build group vector and compute MRCAs
unique_phyla <- unique(na.omit(phylum_vec))
mrca_list <- list()
for (phy in unique_phyla) {
  tips_in_group <- labels[phylum_vec == phy & !is.na(phylum_vec)]
  if (length(tips_in_group) >= 2) {
    mrca_node <- getMRCA(tree, tips_in_group)
    if (!is.null(mrca_node))
      mrca_list[[phy]] <- list(node = mrca_node, n = length(tips_in_group))
  }
}

# 4. Monophyly check
for (phy in names(mrca_list)) {
  tips_in_group <- labels[phylum_vec == phy & !is.na(phylum_vec)]
  if (!is.monophyletic(tree, tips_in_group))
    warning(paste("Non-monophyletic:", phy))
}

# 5. Generate color palette
colors <- setNames(viridis(length(mrca_list)), names(mrca_list))

# 6. Render tree and collapse clades
# Note: mode="mixed" is ggtree default; Rclade forces "max" in circular layouts.
p <- ggtree(tree, ladderize = TRUE)
for (phy in names(mrca_list)) {
  tc <- mrca_list[[phy]]$n
  if (tc > 1) p <- ggtree::scaleClade(p, node = mrca_list[[phy]]$node,
                                      scale = 1 / tc, vertical_only = TRUE)
```

```r
}
for (phy in names(mrca_list)) {
  p <- ggtree::collapse(p, node = mrca_list[[phy]]$node, mode = "mixed",
                        fill = adjustcolor(colors[phy], alpha.f = 0.3),
                        color = colors[phy])
}

# 7. Legend
p <- p + scale_fill_manual(name = "Phylum",
  values = setNames(adjustcolor(colors, alpha.f = 0.3), names(colors)),
  guide = guide_legend(override.aes = list(alpha = 1)))

# 8. Geological timescale
p <- ggtree::revts(p)
p <- p + deeptime::coord_geo(
  dat = list("eons", "eras"), pos = list("bottom", "bottom"),
  xlim = c(-max(node.depth.edgelength(tree)), 0),
  ylim = c(0, Ntip(tree) + 1),
  neg = TRUE, abbrv = TRUE, height = grid::unit(1.5, "line"))

# 9. Theme and save
p <- p + theme(legend.position = "bottom",
               legend.title = element_text(face = "bold"),
               plot.margin = margin(10, 10, 5, 10, "pt")) +
  labs(x = "Time (Ma)")
ggsave("output_baseline.pdf", p, width = 14, height = 10)
```
